# NMJ-Analyser: high-throughput morphological screening of neuromuscular junctions identifies subtle changes in mouse neuromuscular disease models

**DOI:** 10.1101/2020.09.24.293886

**Authors:** Alan Mejia Maza, Seth Jarvis, Weaverly Colleen Lee, Thomas J. Cunningham, Giampietro Schiavo, Maria Secrier, Pietro Fratta, James N. Sleigh, Carole H. Sudre, Elizabeth M.C. Fisher

**Affiliations:** Department of Neuromuscular Diseases, UCL Queen Square Institute of Neurology, University College London, London WC1N 3BG, UK; Mammalian Genetics Unit, MRC Harwell Institute, Oxfordshire, OX11 0RD, UK; UK Dementia Research Institute, University College London, London WC1E 6BT, UK; Department of Genetics, Evolution and Environment, UCL Genetic Institute, University College London, London WC1E 6BT, UK; MRC Unit for Lifelong Health and Ageing, Department of Population Science and Experimental Medicine, University College London, London WC1E 6BT, UK; Centre for Medical Image Computing, Department of Computer Science, University College London, London WC1E 6BT, UK; School of Biomedical Engineering and Imaging Sciences, King’s College London, London W2CR 2LS, UK

## Abstract

The neuromuscular junction (NMJ) is the peripheral synapse formed between a motor neuron axon terminal and a muscle fibre. NMJs are thought to be the primary site of peripheral pathology in many neuromuscular diseases, but innervation/denervation status is often assessed qualitatively with poor systematic criteria across studies, and separately from 3D morphological structure. Here, we describe the development of ‘NMJ-Analyser’, to comprehensively screen the morphology of NMJs and their corresponding innervation status automatically. NMJ-Analyser generates 29 biologically relevant features to quantitatively define healthy and aberrant neuromuscular synapses and applies machine learning to diagnose NMJ degeneration. We validated this framework in longitudinal analyses of wildtype mice, as well as in four different neuromuscular disease models: three for amyotrophic lateral sclerosis (ALS) and one for peripheral neuropathy. We showed that structural changes at the NMJ initially occur in the nerve terminal of mutant TDP43 and FUS ALS models. Using a machine learning algorithm, healthy and aberrant neuromuscular synapses are identified with 95% accuracy, with 88% sensitivity and 97% specificity. Our results validate NMJ-Analyser as a robust platform for systematic and structural screening of NMJs, and pave the way for transferrable, and cross-comparison and high-throughput studies in neuromuscular diseases.

## Introduction

The neuromuscular junction (NMJ) is the peripheral synapse formed by the (a) nerve terminal of a motor neuron, (b) motor endplate of a muscle fibre and (c) ensheathing terminal Schwann cells ^1^. In health, NMJs are dynamic structures with constant remodelling, and continuous innervation/denervation ^1,2^. A permanent morphological change in the NMJ may precede an irreversible denervation process, thought to be the earliest sign of degeneration in multiple human neuromuscular disorders, as well as part of the natural aging process ^3–12^.

Amyotrophic lateral sclerosis (ALS) and Charcot-Marie-Tooth (CMT) disease are devastating neuromuscular disorders. ALS is characterised by progressive degeneration of motor neurons in the brain and spinal cord, leading to skeletal muscle wasting and ultimately death within 2-5 years post symptom onset ^13^. Mutations in at least 30 genes, including *SOD1*, *FUS*, and *TARDBP*, cause ALS ^14–18^. CMT is the most common inherited neurological condition, resulting from mutations in >100 genes, and represents a diverse group of neuropathies affecting peripheral motor and sensory nerves ^19,20^. CMT type 2D (CMT2D) is a subtype caused by mutations in the *GARS* gene ^21^.

Although NMJ pathology is well-known in mouse models of ALS and CMT2D ^6,7,10,22–26^, it is unknown whether pre- and post-synapses degenerate synchronously and which are the earliest structural changes. ALS and CMT2D are distinct diseases, but the underlying peripheral pathology in the neuromuscular system can be studied using similar approaches. NMJs in mice are examined routinely by manual or automatic methods ^27–29^. The manual method entails visual assessment of innervation status (i.e. ‘fully innervated’, ‘partially innervated’ or ‘denervated’), and criteria for NMJ classification can differ, leading to inter-laboratory and inter-rater variabilities, thus limiting further analysis ^30–37^. Automatic methods include NMJ-morph, an innovative approach based on Image-J software, which has been used to analyse NMJ morphology in different species ^38,39^. This platform has widespread use because of its efficiency in measuring morphological NMJ features. However, NMJ-morph uses maximum intensity projection images to capture relevant characteristics of *en-face* NMJs, which may leave behind the great majority of NMJs that acquire complex 3D shapes when wrapping the muscle fibres.

Furthermore, a critical aspect of image analysis lies in cross-study comparison of experiments performed at different times. This can be expedited by using the same thresholding process during analyses and acquiring comparable mean fluorescence intensities (MFI) for wildtype samples. However, in current automatic methods, thresholding is performed manually and sometimes relatively arbitrarily (based on staining observed visually), and MFI and innervation status are not considered. Since change in NMJ innervation status is a primary sign of distal dysfunction in multiple neuromuscular diseases, a systematic, bias-free automatic method is urgently needed to accurately and objectively quantify NMJ characteristics, thereby increasing our understanding of cellular and molecular pathogenesis ^3,6,40^.

To address this need, we developed “NMJ-Analyser”, a robust and sensitive automatic method to comprehensively analyse NMJs. NMJ-Analyser enables quantitative assessment of the native 3D conformation of NMJs using an automatic thresholding method, to accurately capture their morphological features. NMJ-Analyser also incorporates a Random Forrest machine learning algorithm to systematically determine the binary characteristics (‘healthy’ or ‘degenerating’) of NMJ innervation status. This is an innovative approach for large-scale, automated, quantifiable and comprehensive NMJ studies. Thus, NMJ-Analyser associates the two most biologically relevant aspects of the neuromuscular synapse: NMJ innervation with a systematic method to capture topological features.

Here, we describe a comprehensive study of the NMJ using NMJ-Analyser: we evaluated 29 biologically relevant morphological parameters and used a Random Forest machine learning algorithm to diagnose NMJ innervation status. We assessed the morphology of mature NMJs in wildtype mice of two different genetic backgrounds and observed high variability in structure of the endplate (i.e. the post-synaptic acetylcholine receptor clusters), whereas nerve terminals appeared conserved. We went on to evaluate three mouse models of ALS (*SOD1^G93A/+^*, *FUS^Δ^*^14^*^/+^* and *TDP43^M323K/M323K^*) and a CMT2D model (*Gars^C201R/+^*) ^25,26,41,42^. These mice were maintained on different genetic backgrounds, sampled at different ages and disease states, thus giving an ideal opportunity to test NMJ-Analyser. We found NMJ-Analyser was, for the first time, able to detect early structural changes, preceding loss of innervation, in the motor nerve terminals of *FUS^Δ^*^14^*^/+^* and *TDP43^M323K/ M323K^* mice. By comparing our method to NMJ-Morph, we show that NMJ-Analyser detects changes at the NMJ with higher sensitivity.

Our results validate NMJ-Analyser as a robust platform for systematic NMJ analysis, and also shed new light on pathology in the assessed mouse models. Our findings show the value of computer-based approaches for reliable and comprehensive analysis of NMJs to identify early changes, and therefore to potentially aid development of therapeutic interventions in patients with neuromuscular diseases.

## Results

### ‘NMJ-Analyser’ for comprehensive and systematic assessment of NMJs in mice

NMJs are dynamic structures with specific age-related morphological features throughout life ^1,4,43–45^. The morphology of mouse NMJs transitions from poly-innervated (immature, <1-month old mice), mono-innervated (mature, 1 to 18-month old mice) to fragmented/degenerated in aging mice (≥18-month old mice) ^43,45,46^. NMJ morphological studies have been performed qualitatively or semi-quantitively with different methodologies ^6,8,10,24,38,46^; results differ, probably due to the muscles studied, age/genetic background of mice and methods used^1,4,38,44–47^.

In addition, technical variation and differences in quality of staining across samples contribute to variability (batch effect). Thus, a robust and transferrable platform to study NMJs should ideally incorporate a normalization method for reliable systematic analysis. A strength of NMJ-Analyser lies in its ability to normalize parameters across multiple experiments, making cross-study comparison reliable. For example, z-stack images captured at different resolution can be compared as pixel size input in NMJ-Analyser rather than magnification. Additionally, NMJ-Analyser considers a minimum and maximum size for the pre-or post-synaptic NMJ component, reducing the potential error when quantifying staining. Here, we used the same thresholding cut-off across all experiments leading to uniform capture of staining and reduction of background noise.

Here, we present NMJ-Analyser, which consists of four steps to comprehensively study NMJs (Fig.1).

**Fig.1.**
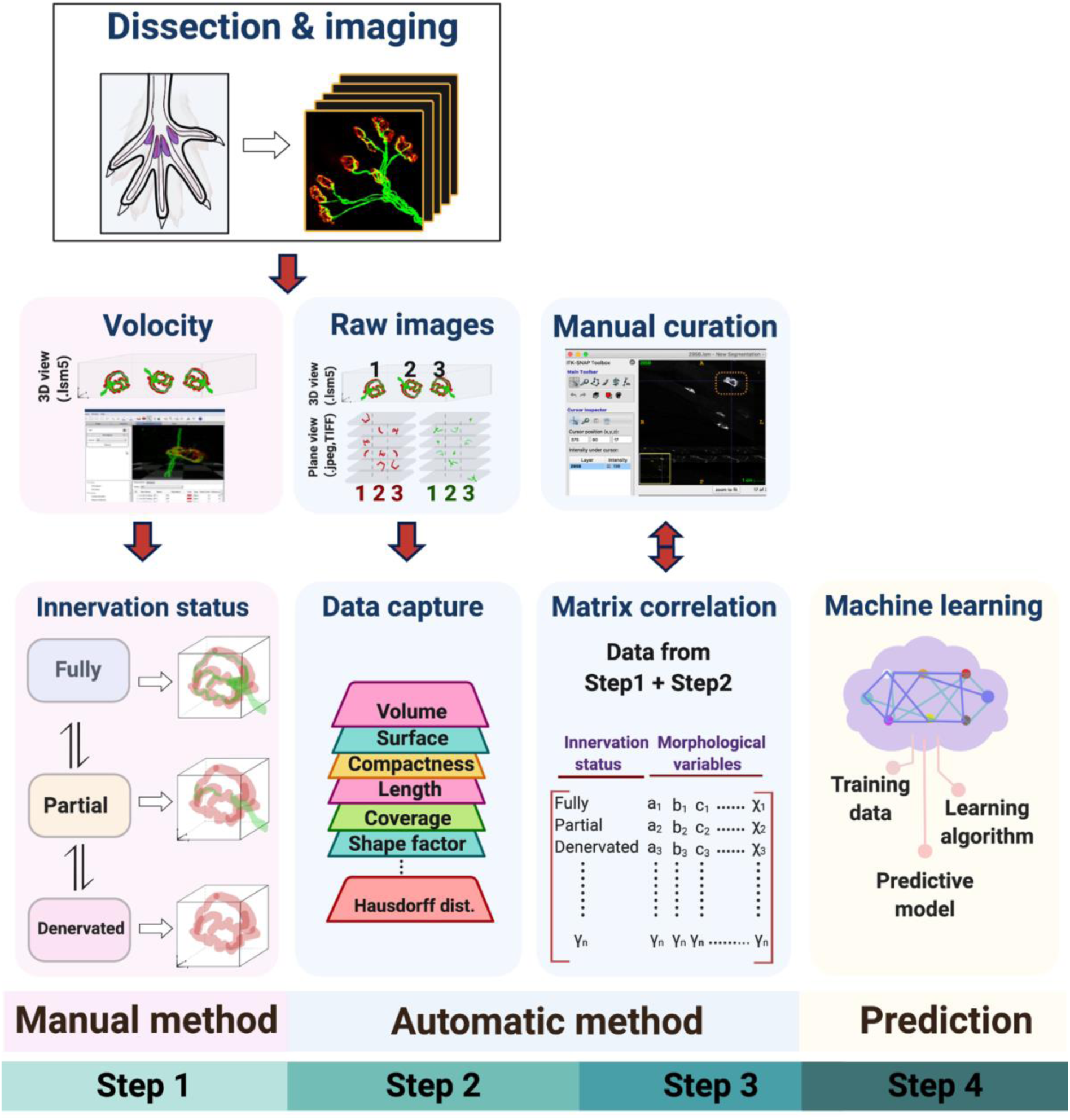
NMJ-Analyser workflow. NMJ-Analyser consists of four sequential steps. (Step 1) Muscles are dissected, immunostained and digitalised (here, hindlimb lumbrical muscles). NMJ innervation status is manually assessed (full, partial, denervated) using an imaging package. (Step 2) Morphological analysis of NMJs requires raw images (plane view) to be formatted (.TIFF/.JPEG/.PNG). Features are captured for individual pre- or post-synaptic NMJs and for parameters involving interactions between these two components. (Step 3) Data from NMJ innervation status and morphological features are integrated into a matrix. Additional manual curation can be performed to confirm the whole individual NMJ was captured and its entire 3D structure analysed using NMJ-Analyser. (Step 4) Curated data output are divided into training and predictive sets. The training set is input into a machine learning model while the predictive set is used to predict the NMJ innervation status. Additional File 2 contains a detailed tutorial of how to use NMJ-Analyser.

#### Step 1: Dissection, staining digitalization and manual assessment of NMJ status

After muscle dissection and staining, NMJ structures are digitalized. NMJ-Analyser requires images in 3D z-stack (3D view) to identify NMJ innervation status. NMJs are classified as fully innervated (‘Full’), partially innervated (‘Partial’) or ‘Denervated’.

#### Step 2: Application of NMJ-Analyser to the stacked raw images and extraction of features

Stacked raw images (plane view) are used to capture the structural features of NMJs using a script we developed in Python. NMJ-Analyser generates twelve biologically relevant parameters for each pre- and post-synaptic structure and five for the interaction between these two NMJ components (29 in total, Table 2). These morphological parameters are non-redundant and biologically relevant.

#### Step 3: Matching manual and automatic assessment

Create a matrix to correlate the *qualitative* information from step 1 and *quantitative* data from step 2, for each NMJ, which allows us to correlate the NMJ innervation status to the respective structural features.

#### Step 4: Training with the Random Forest algorithm and evaluation of testing set

Divide the dataset in the matrix into (i) training data (∼80% of the data) and (ii) testing data (∼20% of the data). The training data are used as an input for a Random Forest machine learning algorithm. The testing data are analysed by the machine learning algorithm to diagnose the innervation status of the NMJs within this dataset. The workflow of NMJ-Analyser can be found in Fig.1 and details about the pipeline are in Additional File 1: Fig.S1.

### Early-mature NMJs display morphological differences compared to mid- and late-mature synaptic structures

Studies of NMJ pathology in small whole-mounted muscles have considerable advantages over analyses in large muscles; such small, thin, flat muscles include lumbricals, FDB and levator auris longus (LAL) ^28,48–51^. These thin muscles do not need sectioning, reducing the variability in antibody penetration, and permit the entire NMJ innervation/denervation pattern to be accurately assessed ^28,48–51^. Furthermore, NMJs in regions with different susceptibility to degeneration can be anatomically defined in small muscles ^12^.

NMJ architecture acquired at early maturity (mono-innervation, pretzel-like shape, 1 month of age mice) is maintained throughout life ^4,45^. However, subtle morphological changes may evade detection as most examinations have been qualitative ^44,45^. Therefore, to quantitively assess such changes, we evaluated biologically relevant NMJ morphological parameters in whole-mounted hindlimb lumbrical muscles in C57BL/6J WT male mice of 1, 3 and 12 months of age (1-month NMJ=early-mature, 3-month NMJ=mid-mature, 12-month NMJ=late-mature; Table 1-2, Fig.2) and C57BL/6J-SJL WT male mice of 1 and 3.5 months of age (Additional File 1: Fig. S2).

**Fig.2.**
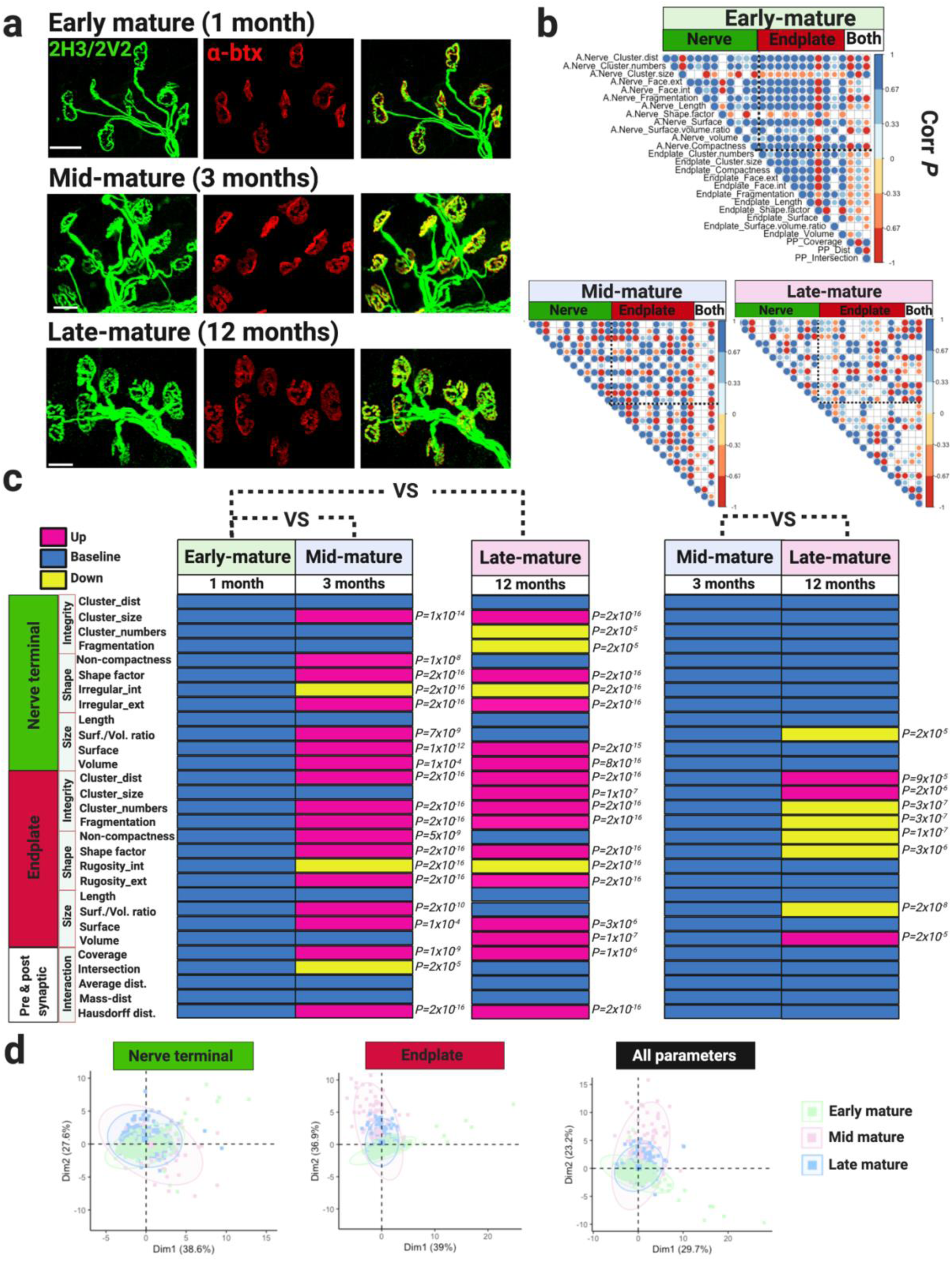
Structural features of healthy NMJs in male C57BL/6J mice. (a) Confocal images of NMJs visualized using nerve terminal markers (2H3+SV2, green) and motor endplate staining (alpha bungarotoxin, α-btx, red). Early-, mid- and late-mature NMJs display mono-innervated axonal input and pretzel-like shape. (b) Matrix correlation plot of nerve terminal and motor endplate parameters, and interaction between them measured in early-, mid- and late-mature NMJs. Positive and negative correlations are given by blue and red scales, respectively (*P*-value ≤ 5×10^-2^). Datapoint showed non-normal distribution and spearman correlation tests were used. Empty circles indicate no significant *P*-value was observed. (c) Module displaying NMJ morphological parameters measured in early-, mid- and late-mature NMJs. Early-mature NMJs are considered as baseline values. Morphological parameters were analysed using Wilcoxon signed-rank test (d) PCA plots displaying variation between early-, mid- and late-mature NMJs. Scale bar = 20 μm.

**Table 1.**
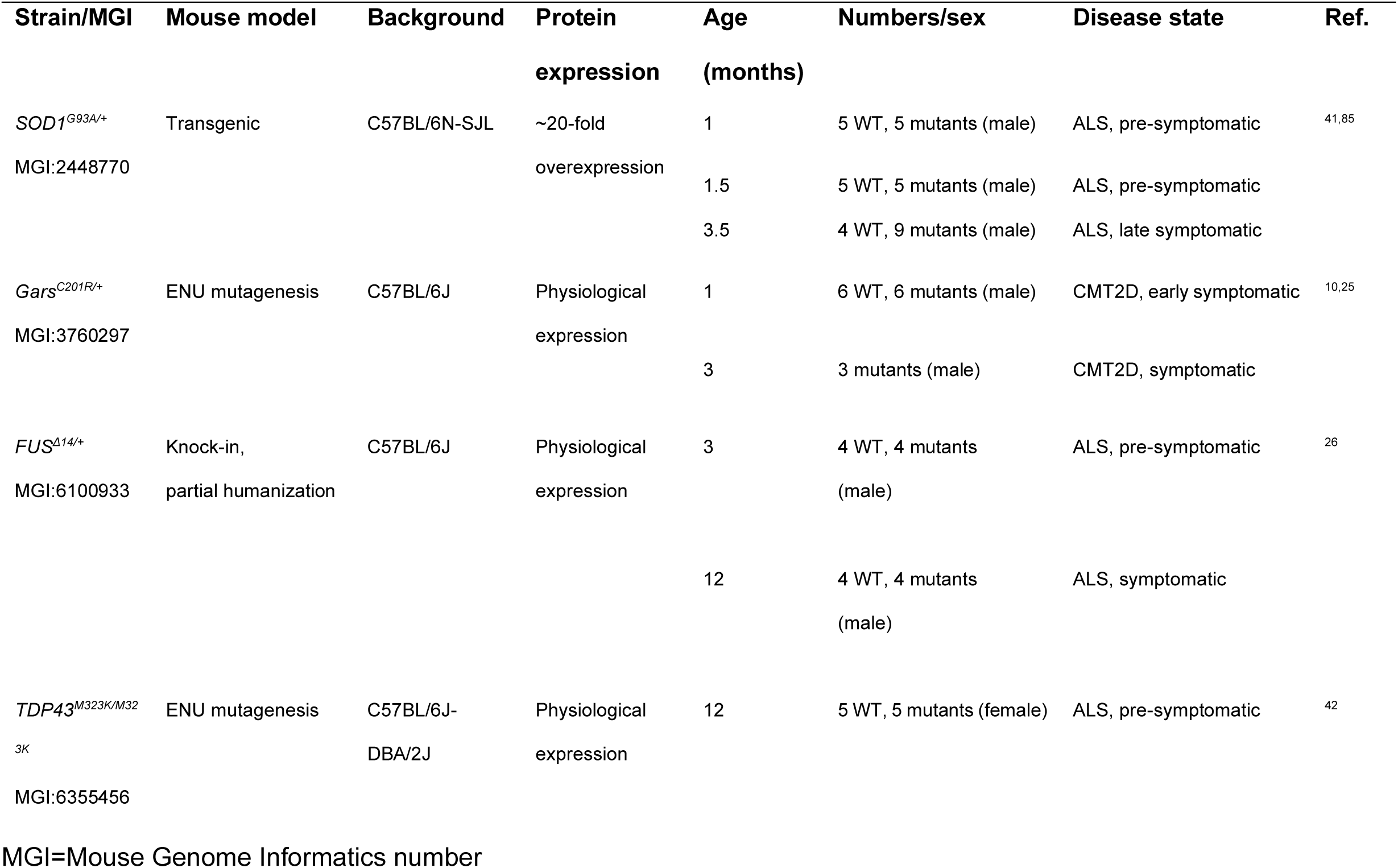
Summary of mutant mice.

Immunolabeling showed conserved NMJ architecture in C57BL/6J mice of 1, 3 and 12 months of age (Fig.2a). Across these timepoints, NMJs were mono-innervated with a pretzel-like shape indicating maturity (Fig.2a). NMJ-Analyser evaluated twelve morphological features of nerve terminals and endplates, and 5 parameters of the interaction between them in these mice (Fig.2b-c). Overall, matrix correlation plot showed that 90% of parameters were significant correlated (71% positive and 19% negative, Additional File 1: Fig.2b, Spearman correlation). Conversely, mid-mature NMJs displayed fewer (74%) significant positive (45%) and negative (29%) correlations (Fig.2b, Spearman correlation). Late-mature NMJs had even fewer (63%) significant correlations between the parameters (42% positive, 21% negative, Fig.2b, Spearman correlation). These results indicate a higher degree of relationship between morphological parameters of early-mature NMJ than mid- or late-mature neuromuscular synapses of C57BL/6J male mice. Similarly, early-mature NMJs from 1-month old C57BL/6J-SJL mice, showed a greater degree of significant correlation between their morphological parameters than those at 3.5 months of age (75% vs 67%, Additional File 1: Fig.S2, Spearman correlation), but this correlation was weaker compared to the one observed in C57BL/6J male mice.

Then, we investigated which parameters are preserved or changed during NMJ maturation. Here, morphological parameters were broadly grouped into ‘integrity, ‘shape’, ‘size’ and ‘interaction’ features. ‘integrity referred to the clusterization characteristics of endplate and nerve terminals, ‘shape’ quantified the overall topology, ‘size’ calculated the dimensions of both NMJ components and ‘interaction’ measured the interaction features of pre- and post-synaptic structures (Table 2, Fig.2). Early-mature nerve terminals had the smallest volume, surface and shape factor, but length remained constant across maturity (Fig.2c). Mid- and late-mature nerve terminals were bigger and presented more significant differences from early-mature pre-synaptic NMJs than between themselves (Fig.2c). Motor endplates showed variability across age with no clear pattern observed across maturity (Fig.2c). Morphological parameters measuring the interaction between nerve terminal and endplate showed that mid- and late-mature NMJ structures were conserved and vary significantly from early-mature architecture (Fig.2c). These results were similar in the C57BL/6-SJL mice (Additional File 1: Fig.S2).

**Table 2.**
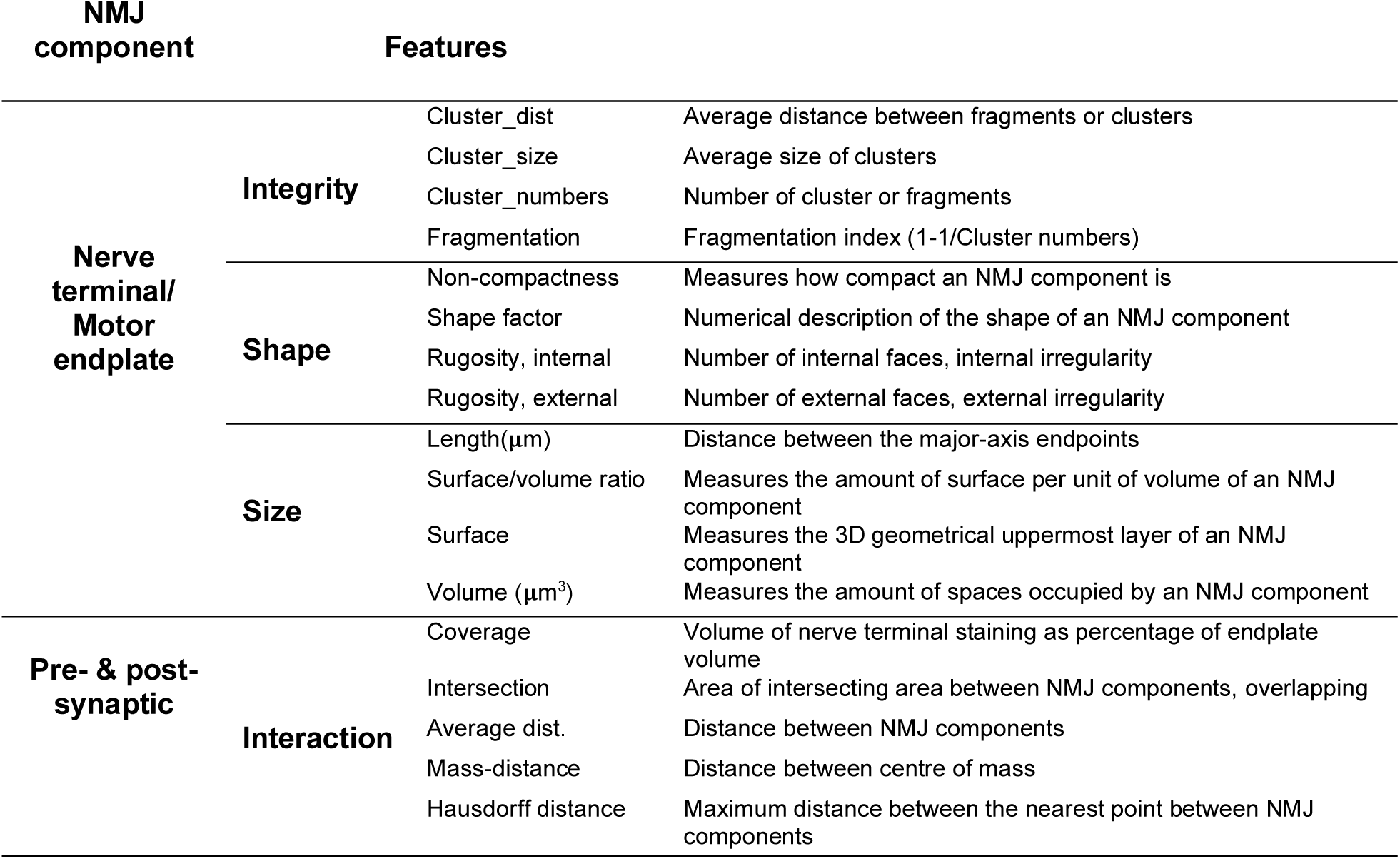
Overview of NMJ morphological features.

| NMJ component | Features |  |  |
| --- | --- | --- | --- |
| **Nerve terminal/**<br>**Motor endplate** | **Integrity** | Cluster_dist | Average distance between fragments or clusters |
|  |  | Cluster_size | Average size of clusters |
|  |  | Cluster_numbers | Number of cluster or fragments |
|  |  | Fragmentation | Fragmentation index (1-1/Cluster numbers) |
|  | **Shape** | Non-compactness | Measures how compact an NMJ component is |
|  |  | Shape factor | Numerical description of the shape of an NMJ component |
|  |  | Rugosity, internal | Number of internal faces, internal irregularity |
|  |  | Rugosity, external | Number of external faces, external irregularity |
| | **Size** | Length($\mu\text{m}$) | Distance between the major-axis endpoints |
|  |  | Surface/volume ratio | Measures the amount of surface per unit of volume of an NMJ component |
|  |  | Surface | Measures the 3D geometrical uppermost layer of an NMJ component |
| | | Volume ($\mu\text{m}^3$) | Measures the amount of spaces occupied by an NMJ component |
| **Pre- & post-synaptic** | **Interaction** | Coverage | Volume of nerve terminal staining as percentage of endplate volume |
|  |  | Intersection | Area of intersecting area between NMJ components, overlapping |
|  |  | Average dist. | Distance between NMJ components |
|  |  | Mass-distance | Distance between centre of mass |
|  |  | Hausdorff distance | Maximum distance between the nearest point between NMJ components |

We further explored changes in morphological variables across maturity using PCA (Fig.2d). We observed that nerve terminals of early-, mid- and late-mature NMJs of C57BL/6J mice cluster together while endplate populations were roughly diverging across these timepoints (Fig.2d). Thus, at all ages analysed, endplates presented greater variability between timepoints than nerve terminals.

### Changes in morphological features in degenerating NMJs

Denervation at NMJs is thought to be the earliest sign of peripheral pathology in multiple mouse models of neuromuscular disease ^6,52,53^. Thus, we determined NMJ innervation status in the hindlimb lumbricals of *SOD1^G93A/+^*, *FUS^Δ^*^14^*^/+^*, *TDP43^M323K/M323K^* and *Gars^C201R/+^* mouse cohorts at the timepoints given (Fig.3a-b). Manual assessment of NMJ innervation status of pre-symptomatic 1-, 1.5-month old SOD1^G93A/+^ and 3-month old FUS^Δ14/+^ mice showed no significant changes when compared to their WT littermates (Fig.3b). Similarly, mildly affected *FUS^Δ^*^14^*^/+^* and *TDP43^M323K/M323K^* mice at 12 months of age had no significant changes in NMJ innervation status compared to WT littermates (Fig.3b). However, we found a significant decrease in the percentage of fully innervated NMJs in 3.5-month old SOD1^G93A/+^ mice (late symptomatic, 50% vs 97% wildtype controls; Wilcoxon signed-rank test, *P-*value=0.002, two-tailed), and early symptomatic 1-month old *Gars^C201R/+^* mice (82% vs 100% wildtype controls, Wilcoxon signed-rank test, *P-*value=0.005, two-tailed) (Fig.3b).

**Fig.3.**
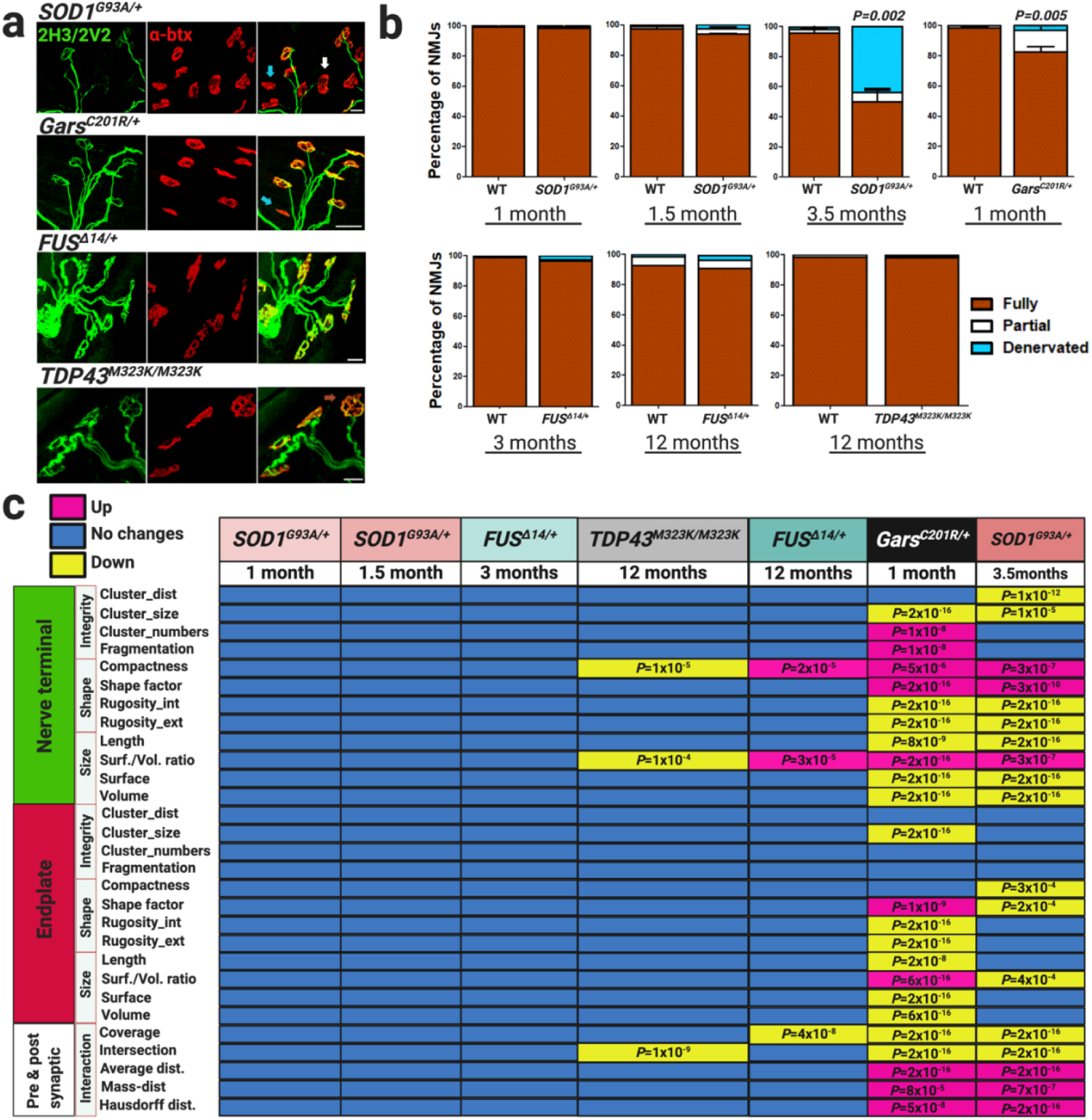
Automatic analysis detects changes before manual assessment of NMJs in ALS and CMT2D mouse models. (a) Confocal images of NMJs visualized using nerve terminal markers (2H3+SV2, green) and motor endplate staining (α-btx, red). Images correspond to *SOD1^G93A/+^* (3.5 months), *Gars^C201R/+^* (1 month), *FUS ^Δ14/+^* (12 months) and *TDP43^M323K/M323K^* (12 months). Arrows point to examples of full (dark orange), partial (white) and denervated NMJs (sky-blue). (b) NMJ innervation status evaluated by eye in whole-mount lumbrical muscles in ALS and CMT2D mouse models. 3.5-month old *SOD1^G93A/+^* and 1-month old *Gars^C201R/+^* mice had a significant reduction in the percentage of fully innervated NMJs (* *P*-value ≤ 0.05, Wilcoxon signed-rank test, two-tailed) (c) Module displaying NMJ morphological parameters measured in ALS and CMT2D strains using NMJ-Analyser. Each timepoint analysed was compared to their respective sex- and aged-matched WT littermates: * *P*-value ≤ 5×10^-4^ was considered significant. Percentage of innervation ± S.E.M. are plotted. Number of mice, WT and mutant: *SOD1^G93A/+^*, 1 month (5+5), 1.5 month (5+5), 3.5 months (4+9); *Gars^C201R/+^* (5+5); *FUS^Δ14/+^*, 3 months (4+4), 12 months (4+4); *TDP43^M323K/M323K^*, 12 months (5+5). Scale bars = 20μm.

Using the same NMJs as above, we investigated structural features in *SOD1^G93A/+^* and *Gars^C201R/+^* mice compared to WT littermates (Fig.3c). *SOD1^G93A/+^* mice had the largest changes in NMJ morphological features (i.e. ‘shape’, ‘integrity and ‘size’ of nerve terminal, endplate); interestingly, these changes were more severe in the nerve terminals than the endplates. 1-month old *Gars^C201R/+^* mice showed extensive and significant variation in volume, length, compactness, surface and length of nerve terminal and endplates; the pre- and post-synaptic components were roughly equally affected (Fig.3c).

A FUS-ALS (*Fus* ^ΔNLS/+^) ^7^ and TDP43-ALS (*TDP43^Q331K/Q331K^*) ^34^ showed reduction in the number of motor endplates in the gastrocnemius compared to WT littermates at 1 and 10 months, and at 10-12 months of age, respectively. To explore whether similar pathology occurs in our mouse strains, we counted the number of endplates per field in all mutant strains and their WT littermates. We found no significant differences in 1- and 1.5-month old *SOD1^G93A/+^* or 3- and 12-month old *FUS^Δ^*^14^*^/+^* strains. However, while 3.5-month old *SOD1^G93A/+^* and 1-month old *Gars^C201R/+^* mice had significant reduction of fully innervated NMJs, motor endplates in these strains were similar to their WT littermates (Fig.3b, Additional File 1: Fig.S3).

### Structural changes in *FUS^Δ^*^14^*^/+^* and *TDP43^M323K/M323K^* mice precede NMJ denervation

Subtle structural changes may be an indication of early NMJ degenerative processes ^1,3,4^. Therefore, we investigated which morphological parameters significantly deviated before NMJ denervation was detectable by eye. We focused on the 12-month old *FUS^Δ^*^14^*^/+^* and *TDP43^M323K/M323K^* strains because, although these mice did not have significant changes in NMJ innervation status, we observed significant alteration of ‘non-compactness’ and ‘volume/surface ratio’ in the pre-synaptic component of both strains, but not in motor endplates (Fig.3b-c). Interestingly, ‘coverage’ and ‘intersection, two parameters defining the interaction between nerve terminal and endplate, were significantly reduced in the *FUS^Δ^*^14^*^/+^* and *TDP43^M323K/M323K^* strains compared to WT littermates, respectively (Fig.3c). Thus, NMJ-Analyser showed the earliest morphological changes in the pre-synaptic nerve in both strains. In summary, NMJ-Analyser detects early and subtle structural changes in NMJs before the denervation process is detectable by eye.

Studies on ALS mouse models have shown that muscle fibre depletion occurs in large hindlimb muscles contributing to the pathology ^23,54–57^. To investigate whether large hindlimb muscles are affected in the FUS^Δ14/+^ strain, we quantified the fibre type composition in fast- and slow-twitch hindlimb muscles (TA, EDL and soleus, Additional File 1: Fig.S4). Fast-twitch TA and EDL muscles in 3- and 12-month old FUS^Δ14/+^ mice showed no variation in fibretype composition nor total number of fibres when compared to WT littermates. Similarly, slow-twitch soleus muscle in 3- and 12-month old FUS^Δ14/+^ mice had no significant differences compared to WT littermates (Fig.S4, Additional File 1: Table S1).

### NMJ-Analyser is more sensitive than NMJ-Morph

NMJ-Morph analyses NMJs that are in an *en-face* position, which occurs when the endplate is completely extended and faces towards the observer (for examples, see Fig.4a i, ii) ^27^. The *en-face* view is ideal to study the size and length of neuromuscular synapses, however, NMJs acquire multiple irregular 3D structures (for example, curved NMJs) as acetylcholine receptors migrate to oppose the motor nerve terminals and the post-synaptic membrane invaginates to enhance neurotransmission efficiency ^45^. Thus, studying only *en-face* NMJs may give an incomplete landscape of structural changes.

**Fig.4.**
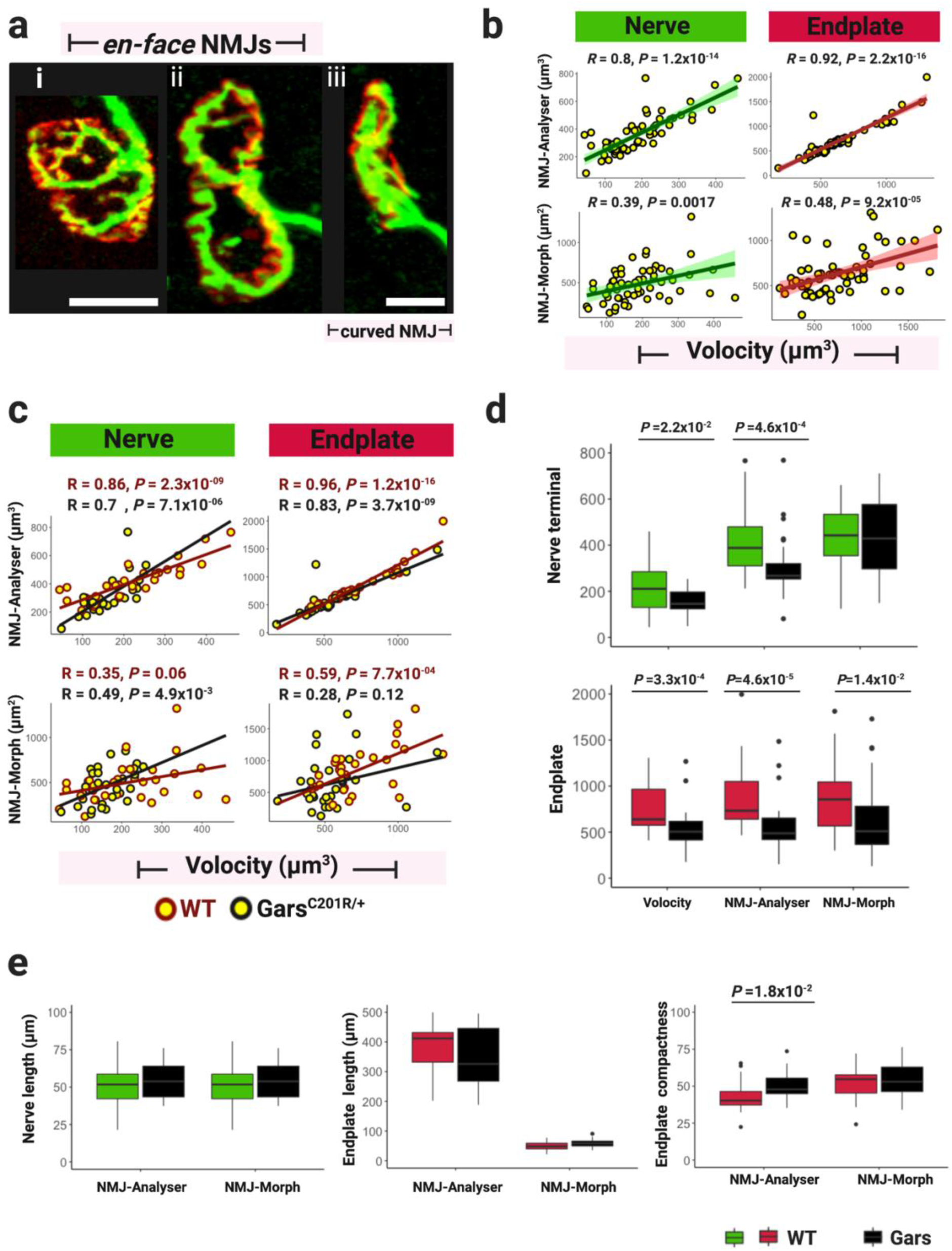
Performance of NMJ-Analyser and NMJ-Morph. (a) Confocal images of (i and ii) *en-face* and (iii) curved NMJs, visualized using nerve terminal markers (2H3+SV2, green) and motor endplate staining (α-btx, red). Images correspond to *Gars^C201R/+^* strain. (b) Correlation of nerve terminal and endplate output obtained using NMJ-Analyser and NMJ-Morph compared to a reference software. (c) Correlation of nerve terminal and endplate output obtained using NMJ-Analyser and NMJ-Morph, divided by genotype. (b-c) Datapoint showed non-normal distribution and spearman correlation tests were used. Green and red shading represent the confidence interval. (d) Boxplot of nerve terminal and endplate output using Volocity, NMJ-Analyser (μm^3^) and NMJ-Morph (μm^2^). (e) Boxplot of nerve terminal and endplate length (μm) and endplate compactness using NMJ-Analyser and NMJ-Morph. c-d outputs were analysed using a using Wilcoxon signed-rank test. Scale bars = 10μm.

Furthermore, NMJ-Morph analyses *en-face* NMJs using maximum intensity Z-stack projections (2D plane). Studying NMJs in their native 3D conformation may give a more complete picture of the ongoing morphological changes, particularly when structural variations are subtle. NMJ-Analyser tackles these drawbacks by analysing the 3D native structure of neuromuscular synapses regardless of their position in a given muscle.

We compared methods analysing NMJs in their 3D native structure versus a 2D maximum intensity projection (NMJ-Analyser vs NMJ-Morph, Fig.4), and compared the outputs using a validated commercial software, Volocity, as a reference. Positions of neuromuscular synapses were identified by an independent evaluator in two different batches: 1-month old *Gars^C201R/+^* (batch 1, Fig.4) and 1.5-month old *SOD1^G93A/+^* mice (batch 2, Additional File 1: Fig.S5) together with sex-, age-matched WT littermates. These batches were selected as the *Gars^C201R/+^* strain had a mild phenotype and *SOD1^G93A/+^* mice presented no NMJ morphological changes at these ages. From all NMJs observed, 28% and 29% were deemed *en-face* to the plane of view in batch1 and batch2, respectively. Thus, imaging *en-face* NMJs may present only a third of the landscape of pathological processes occurring in the lumbrical muscles.

NMJ-Analyser and NMJ-Morph use different approaches to detect NMJ morphological features (i.e. respectively, single vs multiple thresholding, native 3D vs maximum intensity projection), which may lead to different results. To compare the accuracy of NMJ-Morph and NMJ-Analyser outputs, we used Volocity software as a reference. Volocity is a platform to visualize and quantify morphological structures. Thus, the evaluator randomly selected 61 clearly distinguishable *en-face* NMJs from batch 1 and 49 from batch 2. We compared the correlation of nerve terminal and endplate size obtained by NMJ-Analyser (μm^3^) or NMJ-Morph (μm^2^) to Volocity (μm^3^), as a reference value (Fig.4a
). The correlation coefficients of nerve terminal and endplate obtained by NMJ-Analyser were higher compared to NMJ-Morph (*R*=0.8, *P*=1.2×10^-14^ vs *R*=0.39, *P*=1.7×10^-2^ - nerve terminal - and *R*=0.92 *P* =2.2×10^-16^ vs *R*=0.48, *P* =9.2×10^-5^ - endplate -, Fig.4b). Similar results were obtained in batch 2. (Additional File 1: Fig.S5).

We then split batch 1 into WT and *Gars^C201R/+^* NMJs and found that NMJ-Analyser outputs correlated well to Volocity outputs and were more strongly correlated than those of NMJ-Morph to Volocity (Fig.4c). Similar results were obtained in batch 2. (Additional File 1: Fig.S5). We explored whether NMJ-Analyser, NMJ-Morph and Volocity could detect mild structural variations in neuromuscular synapses by comparing nerve terminal and endplate outputs (Fig.4d-e). Nerve terminal and endplate volumes of *Gars^C201R/+^* showed significant reductions compared to WT littermates, using NMJ-Analyser and Volocity (Fig.4d). However, NMJ-Morph output detected significant decreases only in the endplate area of *Gars^C201R/+^* mice, but not in the nerve terminal (Fig.4d). Similarly, we found that endplate compactness was significantly increased using NMJ-Analyser, but not by NMJ-Morph. These results indicate that NMJ-Analyser replicates the findings of Volocity and detected subtle morphological changes in NMJs, while NMJ-Morph did not detect all of the same differences.

### Machine learning diagnoses healthy and degenerating NMJs

Manual NMJ assessment is performed differently between laboratories, impacting cross-comparison studies on neuromuscular innervation ^7,28,30,32,34,35,37,58–60^. Manual assessment had limitations when for NMJ metanalysis. Machine learning is an objective technique that can address this limitation; thus, we used the Random Forest (RF) machine learning algorithm to automatically diagnose healthy and degenerating NMJs (Fig.5).

**Fig.5.**
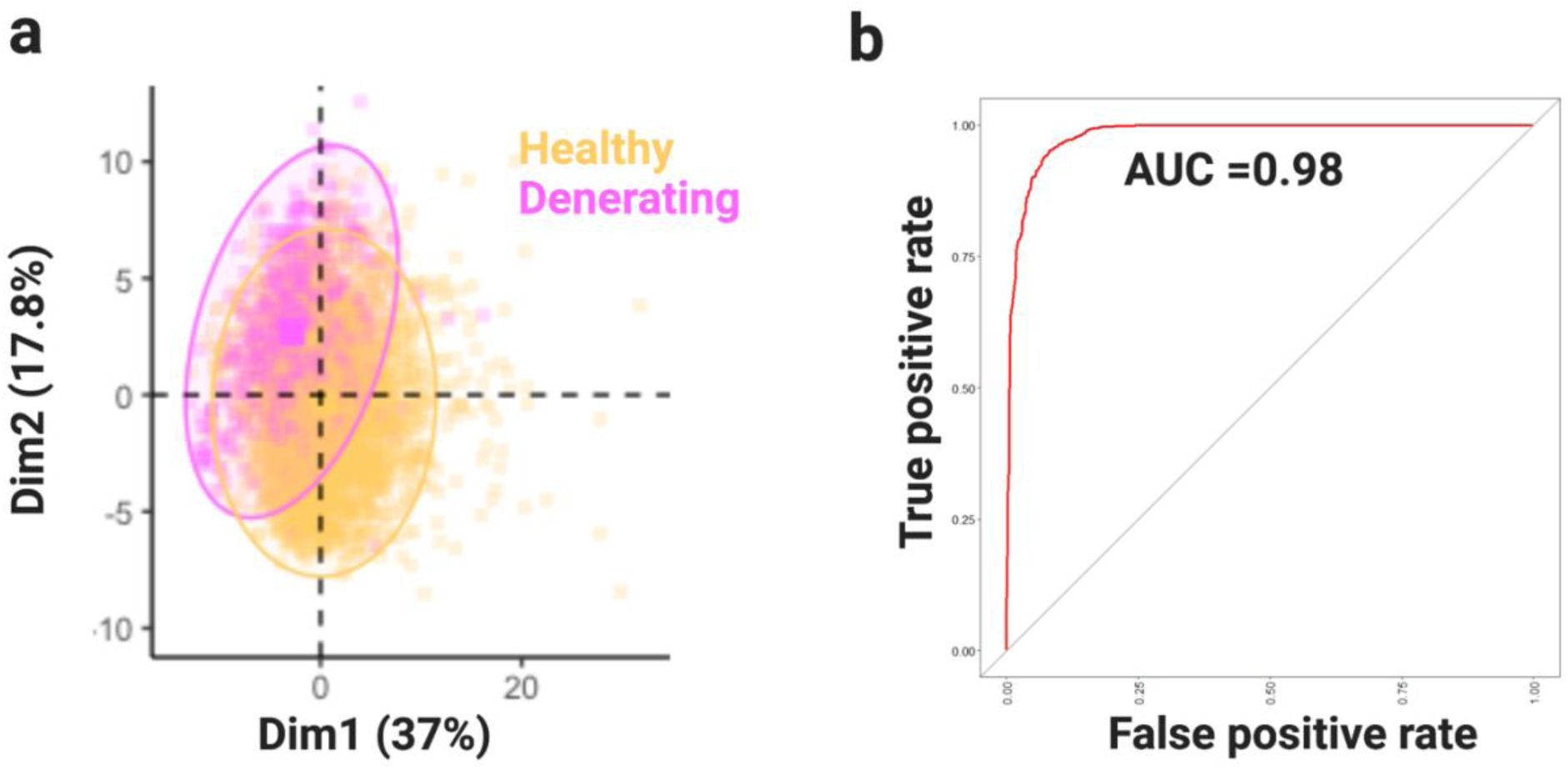
Machine learning algorithm accurately diagnoses NMJ changing. (a) PCA plot displaying healthy (fully innervated) and degenerating (partially innervated and denervated combined) NMJs. (b) AUC plot illustrating the ability of the Random Forest algorithm to differentiate the between healthy and degenerating NMJs.

In our initial observation using PCA, we observed that NMJs in a state of degeneration (i.e. partially, denervated; degenerating) may have some differences from fully innervated synapses (healthy, Fig.5a). Then, we decided to input the morphological features captured to the training RF machine learning algorithm (training dataset). After training step, the RF model diagnosed this binary classification (healthy, degenerating) with 95% accuracy, 88% sensitivity and 97% specificity (Table 3). The area under the curve (AUC), which illustrates the diagnostic ability of our model, was 0.98 (98%, Fig.5b).

**Table 3.** Overview of NMJ diagnosed using machine learning.

| 10-fold RF |  | Morphological features | Type | Overall permutation | Overall Gini |
| --- | --- | --- | --- | --- | --- |
| Accuracy | 0.9552 | Shape factor of endplate | Shape | 100 | 47.2 |
| 95% CI | (0.9417, 0.9664) | Shape factor of nerve | Shape | 58.39 | 18.6 |
| P-value [Acc > NIR] | <2x10<sup>-16</sup> | Intersection | Interaction | 35.1 | 100 |
| Kappa | 0.8604 | Internal rugosity of nerves | Shape | 34.1 | 31.3 |
| Mcnemar's Test (P-value) | 0.6774 | Internal rugosity of endplates | Shape | 31.5 | 17.7 |
| Sensitivity | 0.8809 | External rugosity of nerves | Shape | 27.4 | 22.88 |
| Specificity | 0.9741 | coverage | Interaction | 24.8 | 40.1 |
| Prevalence | 0.2024 | External rugosity of nerves | Shape | 24.1 | 24.7 |
| Balanced Accuracy | 0.9275 | Nerve non-compactness | Shape | 23.3 | 43.2 |
|  |  | Surface/volume ratio of nerves | Size | 21.1 | 24.2 |

We also identified the most important morphological features as ranked by their permutation importance (i.e. the effect removing them from the dataset has on classification accuracy), along with the actual values of permutation importance and Gini importance. Those features belong to the ‘shape’ and ‘interaction’ of NMJ components (Table 3). While more analysis and data from other independent sources is needed to address any possible biases introduced by our sampling, our results do suggest that morphological features can be used to diagnose the healthy/degenerating state of NMJs.

## Discussion

To date, there is no tool for uniform screening of NMJs. Consequently, comparative studies of innervation status and structural analysis are limited, which diminishes our understanding of NMJs in health, ageing and pathology. Here, we describe NMJ-Analyser, a novel platform for morphological screening of NMJs, designed for objective, comparative studies of multiple NMJ structural features. We validated this programme using longitudinal analyses of WT mice, three different genetic ALS mouse models and a CMT2D strain. Using NMJ-Analyser, we have defined changes in WT mice with age and in our pathological models as disease progresses; thus, we have new insights into NMJs in health and in disease, notably including subtle changes in early pathology that may be important for NMJ dysfunction in ALS and CMT2D.

The topology of the NMJ changes from embryogenesis to maturity ^43–45^. Recently, Mech *et al.,* used NMJ-Morph to evaluate five different whole-mounted muscles from 1-month old C57BL/6J WT mice, i.e. early-mature NMJs; muscles analysed were transversus abdominis, flexor digitorum brevis, epitrochleoanconeus and forepaw and hindlimb lumbricals ^61^. Endplate structures in these early-mature NMJs were more variable between muscles compared to their corresponding axon terminals ^61^. NMJ coverage in these muscle (percentage of endplate covered by the axon terminal) in the 1-month old WT mice was ∼63% and the endplate area was ∼2-fold that of nerve terminals ^61^. Data from NMJ-Analyser is similar: 1-month old WT C57BL/6J mice had 66% coverage and endplate volume was ∼1.5-fold that of nerve terminals. Cheng *et al.*, manually assessed *en-face* TA NMJs in female C57BL/6J WT mice from 2 to 14 months of age and reported coverage of ∼72%, which was maintained during the period measured, but endplate fragmentation increased with age ^47^. These results are consistent with our study, in which we found coverage was maintained up to 12 months, but that endplate fragmentation slowly increases with ageing.

We found no differences in individual NMJ-Analyser parameters for the pre-synaptic components i.e. the axon terminals, in WT mice of 3 and 12 months of age. However, both of these timepoints had significant differences from the pre-synaptic component in mice of 1 month of age. Here, as expected, the early-mature pre-synaptic components were significantly smaller than those of mid- (3-month) and late-mature (12-month) NMJs. This could simply be related to age as 3- and 12- month old mice are fully grown while at 1 month they are still developing.

With respect to individual NMJ-Analyser parameters for the post-synaptic components i.e. the motor endplate, in WT mice we found a different non-linear progression of development. Early-mature endplates (1 month of age) had the lowest value parameters, but mid-mature endplates (3 months of age) had the largest parameter values, slightly above those of late-mature (12 months of age) endplates. Thus, endplates appear to follow an inverted “u” pattern, suggesting that the late-mature post-synapse has a more relaxed structure. This difference between mid-mature and late-mature endplates possibly arises from fragmentation and reduction in AChR cluster numbers, which occurs with ageing in healthy WT mice ^47^.

Functional and morphological studies suggest that early decline in muscle function starts at 12-months in C57BL/6J WT mice ^62^. Interestingly, significant reduction in size, orientation, basal calcium content and fission/fusion proteins in mitochondria in the hindlimb FDB muscle were evident in C57BL/6J WT male mice of 10-14 months of age, suggesting incipient muscle decline ^63^. Similarly, sarcopenia and obvious changes in NMJ architecture are evident in 18-month old healthy WT mice ^2,46,64,65^. However, motor unit connectivity, grip strength and muscle contractibility do not vary before 18 months of age ^66^. Hence, it is possible that molecular changes occurring in musculature do not impact muscle function until >18 months of age. More research is needed to clarify why WT nerve terminals appear more stable than endplates, and whether changes within muscle fibres with ageing lead to functional NMJ decline in middle-aged mice (12 months of age).

NMJ degeneration is a process of sequential structural changes occurring in the nerve terminal and/or endplate ^1,67–69^. Two fundamental questions in neuromuscular disease remain unsolved: 1) do the pre- or post-synaptic NMJ components degenerate in a synchronised manner in ALS? and 2) does the skeletal muscle drive NMJ destruction? To validate NMJ-Analyser and to gain new insight into neuromuscular degeneration, we assessed individual NMJ features in a set of mouse models for ALS and CMT2D.

We carried out manual and automatic NMJ analysis in hindlimb lumbrical muscles of *FUS^Δ^*^14^*^/+^* and *TDP43^M323K/M323K^* mice at 12 months of age. Both strains expressed the mutant protein at endogenous levels and had mild yet progressive phenotypes, and are thus ideal models for dissecting early-stage disease processes ^26,42^. We found structural changes in the nerve terminal and the pre- and post-synaptic interaction (for example, ‘compactness’) in both strains, before denervation could be detected by manual analysis, highlighting a major strength of NMJ-Analyser. These results suggest an initial change in neuromuscular morphology, known as remodelling, a process involving the terminal Schwann cells that precedes NMJ dismantling ^2,4,5,67,68,70^.

From our results in the *FUS^Δ1^*^4^*^/+^* and *TDP43^M323K/M323K^* strains, the axon terminal shows initial pathology. Another FUS-ALS model, Ʈ^ON^hFUS^P525L^ knock-in mice of 1 month of age presented strong depletion of synaptic vesicles and mitochondria density in the pre-synaptic portion, but mild reduction in the sarcomere length, and no structural changes in the endplate in the TA muscle, although NMJ denervation was observed ^30^. These findings indicate that initial NMJ remodelling may be associated with alteration of the pre-synaptic morphology and the endplate is affected later.

With more advanced disease progression in FUS-ALS and TDP43-ALS strains, post-synaptic changes i.e. motor endplate pathology, becomes evident. For example, heterozygous 1-month old *FUS^ΔNLS/+^* knock-in mice, which express a truncated FUS protein at endogenous levels, presented reduction in the area and number of endplates in the TA muscles concomitantly with NMJ denervation ^7^. Similarly, transgenic *TD43^M337V/M337V^* mice, which carry a BAC with the full-length human *TARDBP* locus and express the mutant protein at lower levels compared to the endogenous gene, have reduced endplate area at 9 months and NMJ denervation at 12 months of age in the hindlimb lumbrical muscles ^31^. The transgenic mice *TD43^Q331K/Q331K^* of 10-12-month of age had reduction of motor endplates per section and NMJ denervation ^34^.

The slow progressive phenotype of *TDP43^M323K/M323K^* and *FUS^Δ1^*^4^*^/+^* mice did not result in NMJ denervation or reduction in endplate numbers, and fibre typing studies in 3- and 12-month old *FUS^Δ14/+^* mice showed no significant changes. More advanced stages of neuromuscular degeneration present with NMJ denervation, endplate loss and sometimes concomitant muscle fibre changes ^2,7,31,34,68^. Thus, as fibre type remodelling was observed in *TD43^Q3331K/Q331K^* (atrophy fibres, regenerating fibres with centralized nuclei), at 10-12 months of age, it is likely that our 12-month *TDP43^M323K/M323K^* mice were in an early NMJ remodelling process ^34^.

The *SOD1^G93A/+^* mice have an aggressive phenotype because they overexpress the mutant protein ∼24-fold times higher than endogenous gene and thus have a rapid rate of NMJ pathology, which has a preference for fast-twitch muscles ^6,23,40,41,53,68,71^. We analysed manually the NMJ innervation status in the lumbricals of *SOD1^G93A/+^* mice at 1, 1.5 and 3.5 months of age, and the last timepoint showed significant denervation. Our result for the 3.5 month timepoint was consistent with previous reports where ∼50% or more NMJ denervation was found in fast-twitch muscles (medial gastrocnemius, TA and EDL) ^6,22,23,53,72^. However, two independent studies found that *SOD1^G93A/+^* mice had either a small (∼8%, EDL) or prominent (∼50%, gastrocnemius and TA) NMJ denervation at 45 and 47 days of age, respectively. This apparent discrepancy with our results may be explained, in part, by the sex of mice used, muscle fibretype composition, muscle studied and cut-off parameters used for full, partial and denervation NMJ status ^6,60^. Moreover, the methodology of manual NMJ analysis in mouse models varies across laboratories, and different cut-off parameters for NMJ innervation status limit cross-comparison studies ^6,7,10,25,26,30,32–35,36,37,58,59^.

The *Gars^C201R/+^* mice, which model CMT2D, presented a mild denervation phenotype associated with pre-synaptic and post-synaptic structural NMJ changes at 1 month of age: ∼80% of hindpaw lumbrical NMJ were fully innervated, consistent with a recent report ^73^. At this age, the ‘integrity’ parameter, which measures fragmentation and clusterization of nerve terminals and endplates, was increased on average while parameters quantifying size (volume, length, surface) were significantly decreased compared to WT littermates. This is consistent with a previous report that endplate area of 1- and 3-month old *Gars^C201R/+^* mice was 9% and 12% less than WT, respectively, although this did not reach significance ^10^. A combination of delayed NMJ development, aberrant interaction between glycyl-tRNA synthetase (GlyRS) and the tropomyosin kinase receptor (TrK) receptors (some of which are found at the NMJ) and differential muscle involvement, all appear to contribute to the *Gars^C201R/+^* mouse phenotype ^10,73,74^.

In summary, we found that subtle quantifiable structural alterations occur primarily in the nerve terminals in 12-month old *FUS^Δ14/+^* and *TDP43^M323K/M323K^* strains. We also showed *FUS^Δ14/+^* mice had no fibretype switching nor reduction in the numbers of fibres in fast- and slow-twitch muscles at 3 and 12 months of age. Our data suggest that in our FUS and TDP43 ALS physiological models, the pre-synaptic NMJ component degenerates before the endplates and not synchronously. Furthermore, it is unlikely that skeletal muscle drives NMJ destruction as appear to be the case in Spinal-bulbar muscular atrophy disease (SBMA) ^75^.

In the transgenic aggressive ALS model *SOD1^G93A/+^*, pre-synaptic changes were considerably more pronounced than post-synaptic alterations and we did not see reduction in the number of endplates, at the ages sampled, again, suggesting that pre-synaptic changes precede NMJ pathology and skeletal muscle does not drive NMJ denervation.

The mild CMT2D model *Gars^C201R/+^* presented similar changes in pre- and post-synaptic NMJ components, indicating a different pathomechanism from ALS models, perhaps due to the primary site of motor neuron pathology (for example, cell body versus distal synapse).

NMJ-Analyser showed higher sensitivity than manual NMJ methods and NMJ-Morph, and can quantify NMJs in their native 3D structure independently of their shape acquired in the muscle fibres -- this is particularly important because NMJ topology may change during disease ^1,4,40,43^. These morphometric comparisons require capturing the complete density of three-dimensional shape and interrelations, not only distance or ratio, as needed in two-dimensional images ^76–82^. Furthermore, image thresholding, multiple validations of the pipeline and coupling to a machine learning algorithm make our programme a robust and sensitive tool for structural studies of neuromuscular synapses.

NMJ-Analyser is a reliable tool for NMJ topological analysis, but some morphological characteristics were not included in the study. Those parameters include terminal sprouting and poly-innervation and terminal Schwann cell coverage. The code being open-source, any additional contribution of feature is welcome. Another limitation of our study lies in motor endplate quantification: a more precise method is to count the complete number of post-synaptic structures in a given muscle, however, this may technically difficult to achieve.

Compared to the semi-automated analysis tool in NMJ-Morph, NMJ-Analyser showed greater sensitivity and stronger correlation in detecting morphological NMJ parameters in wildtype and mutant mouse strains. Notably, we were able to diagnose NMJ innervation status confidently using machine learning and demonstrated that NMJ-Analyser is an advancement on the widely-used tool NMJ-Morph. To our knowledge, these are the first data demonstrating that structure and innervation state of NMJs can be studied systematically and in an automatic and integrated manner. Our results validate NMJ-Analyser as a unique and robust platform for systematic analysis of NMJs, and pave the way for transferrable and cross-comparison studies in neuromuscular disease.

## Methods

### Animals

Three ALS mouse strains (*SOD1^G93A/+^* ^41^, *FUS^Δ14/+^* ^26^ and *TDP43^M323K/M323K^* ^42^), a CMT2D mouse model (*Gars^C201R/+^*) ^25^ and their corresponding sex- and age-matched wildtype littermates were used to study NMJ pathology (Table 1). All strains, except the transgenic line *SOD1^G93A/+^*, express the mutant protein at physiological levels; details on the genetic background, sex, age and number of mice studied are in Table 1.

All mouse handling and experiments were performed under license from the United Kingdom Home Office in accordance with the Animals (Scientific Procedures) Act (1986) and approved by the University College London – Queen Square Institute of Neurology Ethics Committee.

### Muscle collection and immunohistochemistry

Lumbrical and flexor digitorum brevis (FDB) muscles located in the hindlimb paws of mice were dissected and stained, as previously described ^28,83^. Briefly, after dissection, tissues were fixed with 4% paraformaldehyde (PFA, Sigma-Aldrich) for 8-10 min in phosphate buffered saline (PBS) then permeabilized with 2% PBS-Triton X-100 for 30 min to increase antibody penetration. Then, blocking solution (5% donkey serum in 0.05% PBS-Triton X-100) was added to the tissue for 1 h followed by multiple washes of 15-20 min each. Overnight incubations with mouse anti-synaptic vesicle 2 (SV2, 1/20, DHSB) and mouse anti-neurofilament medium chain (2H3 1/50, DSHB) in blocking solution at 4°C were performed to visualise the pre-synaptic NMJ component. After overnight incubation, multiple washes were conducted with PBS-1X for at least 1 h. AlexaFluor-488 anti-mouse IgG secondary antibody (1/2000, Invitrogen) was added together with labelled α-bungarotoxin (to visualise post-synaptic acetylcholine receptors) in blocking solution for 1.5 h at room temperature (RT) (α-btx, Life Technologies). Then, muscles were washed several times and whole-mounted for imaging.

### NMJ imaging

Images of NMJs were obtained using a Zeiss LSM 710 confocal microscope (Zeiss, Germany). All strains were imaged at 512×512 resolution, except for the *FUS^Δ^*^14^*^/+^* strain, which was at 1024×1024. Z-stack images were decomposed into individual pre- and post-synaptic planes for posterior analysis (.TIFF, .PNG and .JPEG).

### Manual NMJ analysis

Stacks of images containing complete NMJs were visualised in Volocity software (version 6.5, Perkin Elmer) and their innervation status (fully innervated, partially innervated, denervated) was manually assessed, as previously described ^24,28^. When a nerve terminal and motor endplate overlapped with at least 50% coverage the NMJ was considered ‘fully innervated’. When between ≥20% and ≤50% overlap occurred, NMJs were considered ‘partially innervated’. Vacant motor endplates were considered ‘denervated’ NMJs. Only NMJs whose innervation status were clearly defined were considered for this study.

### Automatic NMJ analysis

For each manually assessed NMJ, we obtained the 3D position [*P* (x, y, z)] in Euclidean space using Volocity. This step was performed to identify NMJ position and accurately correlate innervation status with the corresponding morphological characteristics (Fig.1, Additional File 1: Fig.S1). In parallel, the stacked images were used as input to the “NMJ-Analyser” framework and to extract their morphological features for nerve terminal, motor endplate and the interaction between them (Fig.1). 29 features were analysed for each NMJ (Table 2). Manually assessed NMJs were then matched to the automatically extracted output using the minimum Euclidean distance between centre of mass. Since Z-stack images frequently contained superimposed NMJs, we only considered NMJs that were clearly differentiated one from another for analysis (Fig.1, manual curation); on average, NMJs need to be separated by at least 20 µm. The final number of NMJs identified by both manual and automatic analysis was 6,728.

### Validation of NMJ-Analyser and manual curation

To confirm the reliability of NMJ-Analyser, we validated key morphological parameters (volume of nerve terminal and endplate) using Volocity software (Additional file 2: Fig.3). Additionally, NMJ-Analyser automatically provides the centre of mass of individual NMJs (3D position, [*P* (x, y, y)]). Centre of mass refers to the average NMJ position in 3D Euclidean space, which is unique for every synapse. The centre of mass position was used to correlate NMJ innervation status identified by Volocity with paired morphological parameters (Fig.1, matrix correlation). We also used ITK SNAP, a 3D software viewer, to visually confirm that NMJ-Analyser detected the whole individual NMJ structure and captured its entire 3D morphological structure (Fig.1, manual curation) ^84^.

### Machine learning

We aimed to reduce the variability in human interpretation by using machine learning. The supervised machine learning technique Random Forest was used to diagnose the innervation status of NMJs (healthy or degenerating) using automatically extracted morphological characteristics. From the total number of NMJs (Additional File 1: Table S2), the dataset was randomly split into a training dataset consisting of 80% of the full dataset and a testing dataset made up of the remaining 20%. While the ratio of degenerating to healthy NMJs was not deliberately preserved, the two datasets were made up of 20% and 25% degenerating respectively. We also created an equalised dataset taking 1000 rows at random from degenerating and healthy

The training and testing models were trained to optimise accuracy. In our testing, prioritising kappa (caret’s other option for classification) did not make a substantial difference to the outcome. The random forest models were trained using caret’s train function, creating one model using the full training dataset and one using the equalised one. Both of these models had 10-fold cross validation. The innervation of the NMJs comprising their respective test datasets were diagnosed and compared to the actual values.

### NMJ-Morph analysis

Semi-quantitative analysis of *en-face* NMJs was performed as previously described ^27^. Maximum intensity projections from individual pre- and post-synaptic staining were obtained using the black Zen software (Zeiss, Germany). The Image J manual thresholding method was used as it gave the most consistent results across different methods. Then, binary images were obtained after background elimination. We measured area and length of nerve terminals, and acetylcholine receptor (AChR) area, diameter and area of endplate and compactness.

### Motor endplate counts

Motor endplates of lumbrical muscles were counted in 8-12 fields per mouse. Motor endplates from wildtype and mutant littermates were counted at 40x magnification, except 3- and 12-month old *FUS^Δ14/+^* and their wildtype littermates, which were quantified at 20x magnification.

### Fibre type counts

Tibialis anterior (TA), extensor digitorum longus (EDL) and soleus muscles were snap frozen and transversally sectioned at 20 µm thickness using a microtome (Leica, Germany), and left to dry for 1-2 h. Sections were rehydrated with PBS-1X at RT. Non-specific blocking solution (10% donkey serum in 0.01% PBS-Tween 20) was added to the tissue for 1 h, followed by multiple 15-20 min washes. Overnight incubation consisted of blocking buffer with mouse anti-fibre type I (IgG2b,1/10, DHSB); mouse anti-fibre type IIa (IgG1, 1/20, DHSB), mouse anti-fibre type IIb (IgM, 1/20, DHSB) and rabbit anti-dystrophin (1/750, DHSB), which were used to recognize borders of individual fibres. After overnight incubation and multiple washes with PBS-1X for 1 h, we incubated the sections for 2 h with the following secondary antibodies: anti-mouse IgG2b Alexa Fluor-405, anti-mouse IgG1 Alexa Fluor-488, anti-mouse IgM Alexa Fluor-564 and anti-rabbit Alexa Fluor-633. Then, muscle sections were washed several times with PBS and mounted for imaging.

Hindlimb sections were imaged at 20x magnification using a Zeiss 7100 Confocal Microscope (Zeiss, Germany). Images were counted using the ‘CellCounter’ plugin on Image J.

### Statistical analysis

Data analysis and plotting were performed using R/RStudio. Schematic graphics found in Fig.1 and Additional File 1: Fig.S1 were created using BioRender. Different plot types were developed to represent various level of analysis. During the study, a single batch experiment included mutants and sex- and age-matched WT littermates, thus, technical variation between genotypes within a batch was assumed null. When comparing multiple batches (i.e. for C57BL/6J and C57BL/6J-SJL male mice, longitudinal study), the following normalization procedure was performed. Normalization protocol can be found at Additional File 2.

### *P*-value and normality distribution of data

Different approaches to test statistical significance were performed. When comparing NMJ morphological features between genotypes, we tested the normality of data distribution using the Shapiro-Wilk test. In most cases, our data follow a non-normal distribution, thus we conducted non-parametric statistical tests: the Wilcoxon signed-rank test or Kruskal-Wallis test (one-way ANOVA) to compare two or three groups, respectively. *P*-value adjustment was performed when three or more groups (or variables) were compared simultaneously (Benjamin-Hochberg -BH-post-hoc test).

When comparing manual NMJ innervation status, motor endplate counts and fibre type counts, we set significance as *P*-value ≤ 5×10^-2^, as we compared the percentage or average obtained per mouse (biological replicates). To compare the performance of Volocity, NMJ-Analyser and NMJ-Morph, a *P*-value ≤ 5×10^-2^ was considered significant as we used <100 NMJs. When we compared individual NMJs, rather than mice, we used a restricted adjusted *P*-value ≤ 5×10^-4^ as cut-off for statistical significance.

Correlation matrix plots were obtained to measure the relationship between two morphological variables. A *P*-value ≤ 5×10^-2^ was considered significant. Spearman correlation plots were generated using the *corrplot* package in R/RStudio.

## Acknowledgments

We are grateful to Prof. Thomas Gillingwater, Prof. Adrian Issacs, Dr Bernadett Kalmar, Dr J. Barney Bryson and Dr Anny Devoy for their helpful suggestions during development of NMJ-Analyser.

## Author Contributions

A.M.M. and E.M.C.F. conceived the experiments; A.M.M., S.J, W.C.L., J.N.S., and C.H.S. performed the research; T.J.C. provided tissues samples, A.M.M., S.J. and C.H.S. analysed the data; T.J.C., G.S., M.S., J.N.S, E.M.C.F., P.F. and C.H.S. provided expertise and discussion; A.M.M. and E.M.C.F. wrote the manuscript with inputs from all authors. All authors agreed on the submission of this work.

## Additional Information

Supplementary information accompanied this paper can be found at https://github.com/csudre/NMJ_Analyser and https://github.com/SethMagnusJarvis/NMJMachineLearning.

## Additional File 1

**Fig.S1.**
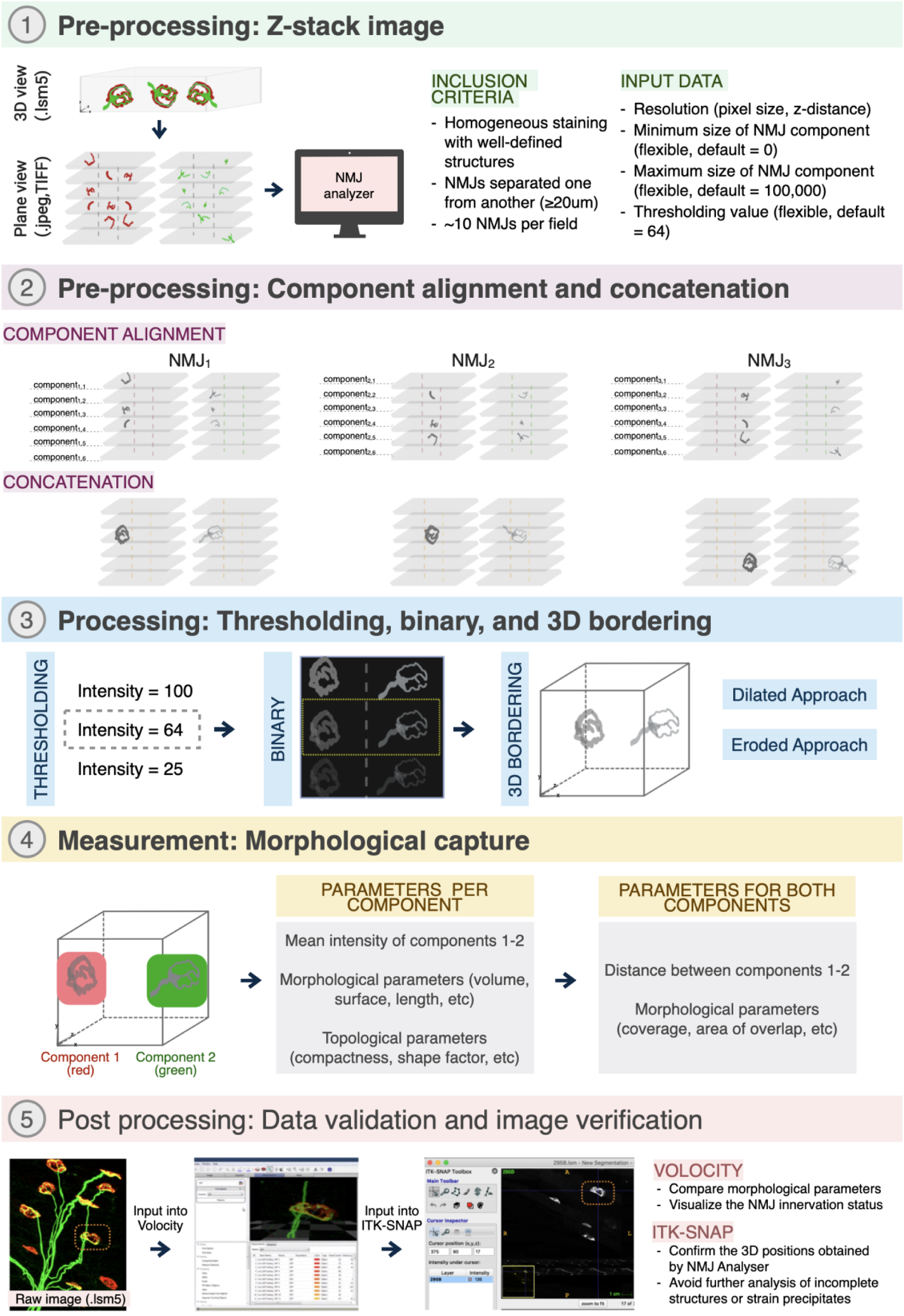
Workflow of image morphological analysis using NMJ-Analyser. Morphological analysis of NMJs requires (1) pre-processing, (2) processing, (3) measurement and (4) post-processing of images. (1) Images are required to be in adequate format for identification of NMJ innervation status and for posterior capture of morphological features (3D and plane images, respectively). (2) Pre-processing step is used for capturing automatic morphological features. Images concatenation is used to assigned a 3D position to each NMJ component (centre of mass, [*P* (x, y, y)]). (3) Processing refers to the cut-off intensity used to define and differentiate the staining from background (thresholding). It is important to keep the same threshold between batches for reproducibility. (4) Measurement refers to the capture of morphological features of pre- and post-synaptic NMJ. (5) Post-processing or manual curation is required as NMJ-Analyser gives a large output automatically. When analysing images in a high throughput manner, antibody precipitates or incomplete NMJs may be identified and their features collected. Thus, using the 3D position of individual objects identified in step 2-4 can be clearly identified using ITK-SNAP viewer (freely available). These antibody precipitates or incomplete NMJs identified can be then easily dropped for further analysis.

**Fig.S2.**
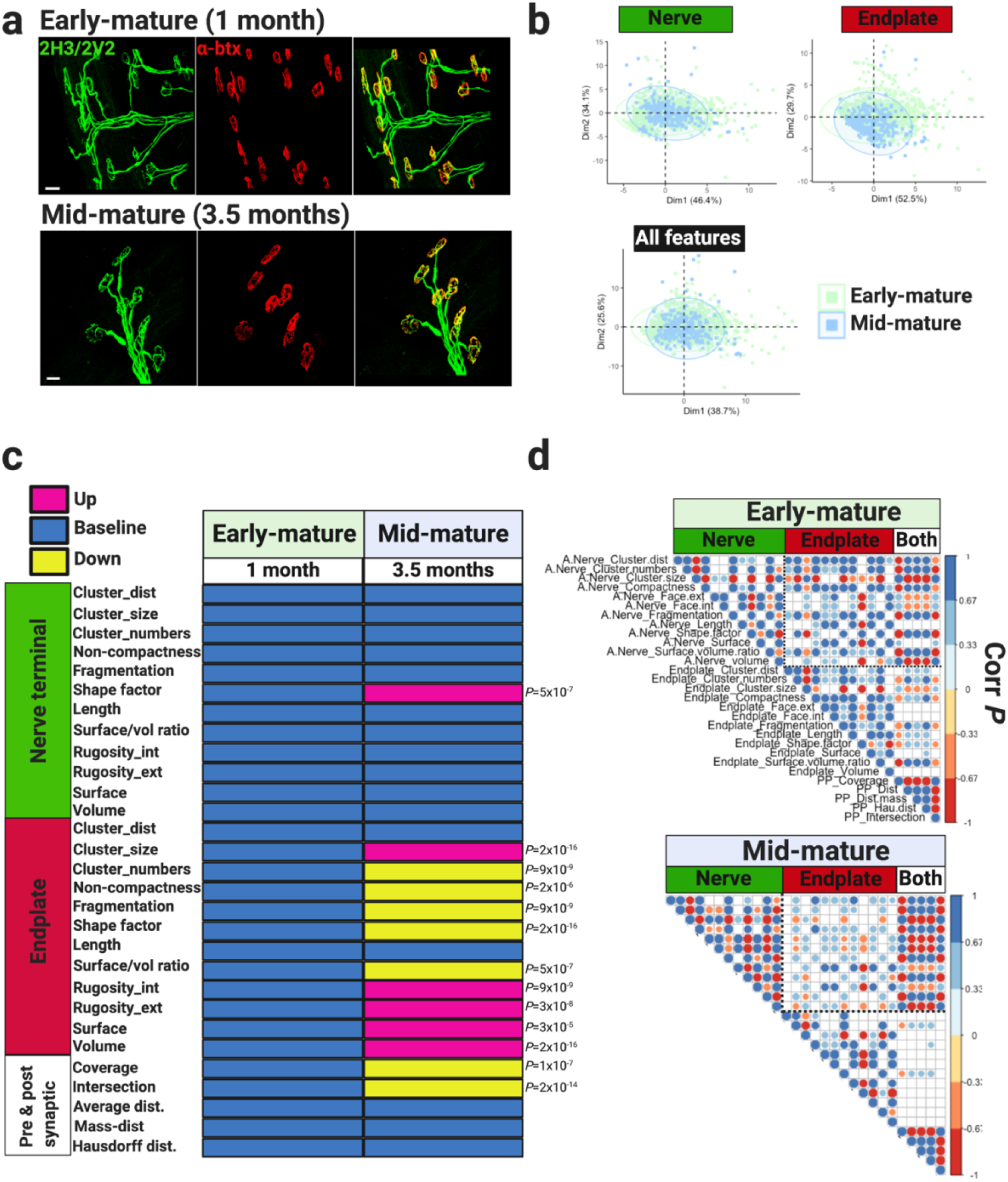
Structural features of healthy NMJs in male C57BL/6J-SJL mice. (a) Confocal images of NMJs visualized using nerve terminal markers (2H3+SV2, green) and motor endplate staining (α-btx, red). Early- and mid-mature NMJs display mono-innervated axonal input and pretzel-like shape. (b) PCA plot showing no major cluster differentiation between early- and mid-mature NMJs. (c) Module displaying NMJ morphological parameters measured in early- and mid-mature NMJs. Early-mature NMJs are considered as baseline values. Morphological parameters were analysed using a using Wilcoxon signed-rank test (d) Matrix correlation plots of nerve terminal and motor endplate parameters, and interaction between them measured in early- and mid-mature NMJs. Positive and negative correlations are given by blue and red scales, respectively (*P*-value ≤ 5×10^-2^). Datapoint showed non-normal distribution and spearman correlation tests were used. Empty circles indicate no significant *P*-value was observed. Number of mice, early- and mid-mature: (5+4). Scale bars = 20μm.

**Fig.S3.**
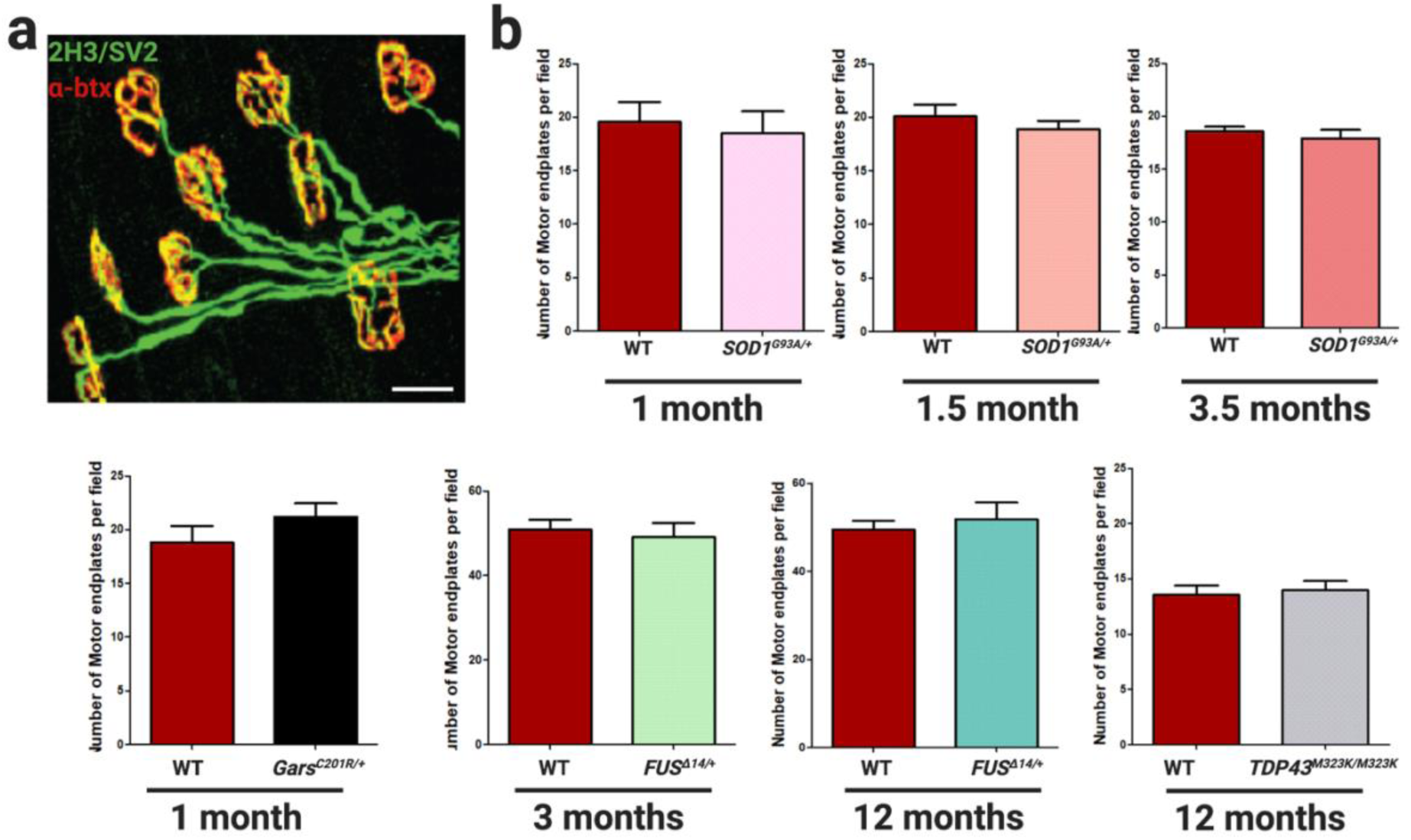
Motor endplate counts in ALS and CMT2D mouse models. (a) Lumbrical NMJs from 1.5-month-old WT mice visualized using nerve terminal markers (2H3+SV2, green) and motor endplate staining (α-btx, red). (b) Bars represent average of motor endplates per field; 8-12 fields per mouse were counted. Number of mice, WT and mutant: *SOD1^G93A/+^*, 1-month (5+5), 1.5-month (5+5), 3.5-months (4+9); *Gars^C201R/+^* (5+5); *FUS^Δ14/+^*, 3-months (4+4), 12-months (4+4); *TDP43^M323K/M323K^*, 12-months (5+5). Mean ± S.E.M were plotted. Scale bar = 20μm.

**Fig.S4.**
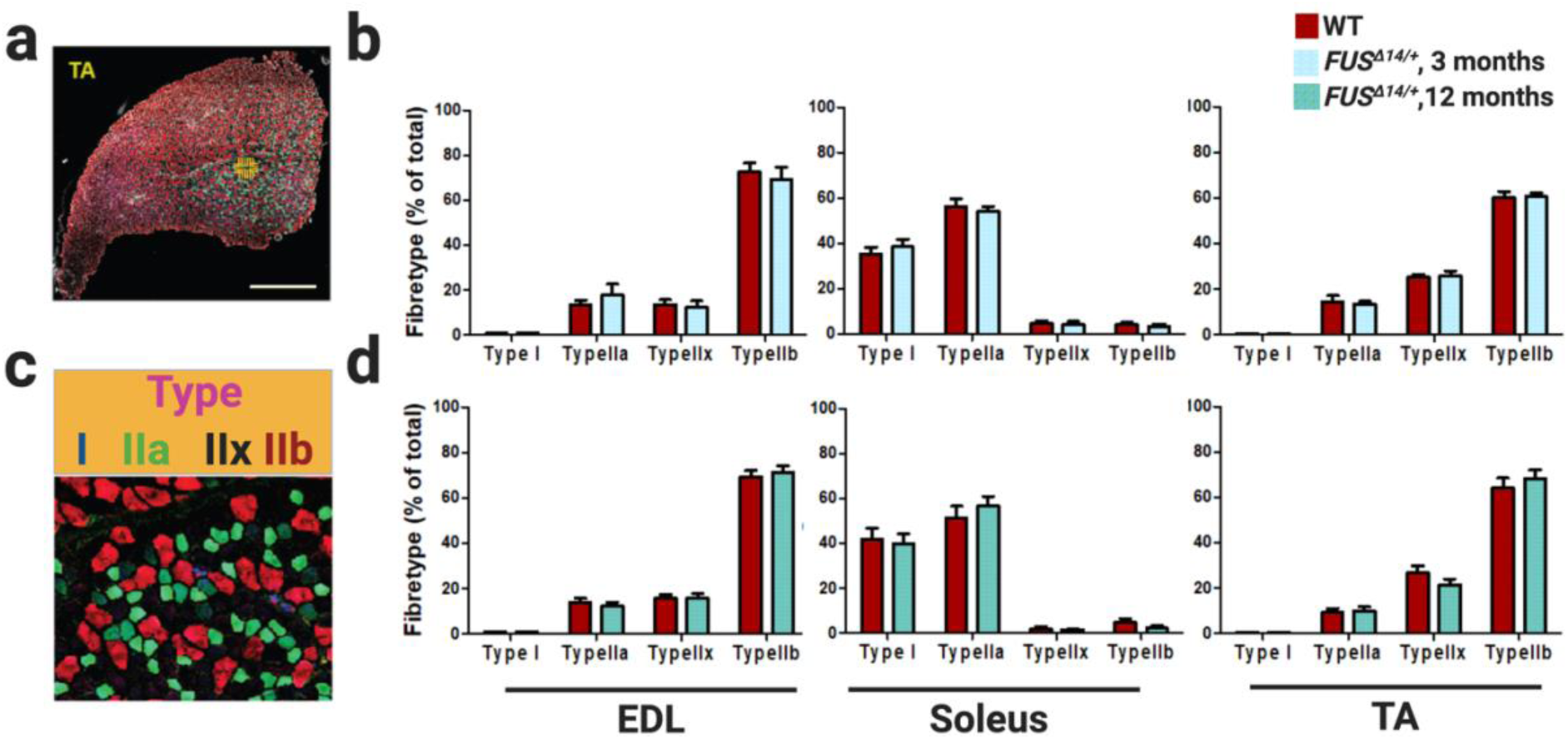
Fibretype composition in hindlimb muscles of *FUS^Δ14/+^* mice. (a) Transversal section of TA muscle visualized using fibre markers type I, IIa and IIb. (b) Fibretype composition of EDL, soleus and TA hindlimb muscles in 3-month old WT and FUS^Δ14/+^ littermates. (c) High-magnification of TA shown in (a), displaying individual fibretypes. Fibretype I (blue), IIa (green), IIx (negative staining) and IIb (red). (d) Fibretype composition of EDL, soleus and TA hindlimb muscles in 12-month old WT and *FUS^Δ14/+^* littermates. Number of mice, WT and *FUS^Δ14/+^*: 3-months (7+8), 12-months (7+8). Male and female mice at each timepoint (3+4 or 4+4). No significant sex-differences were observed. Mean ± S.E.M were plotted. Scale bar = 20μm.

**Fig.S5.**
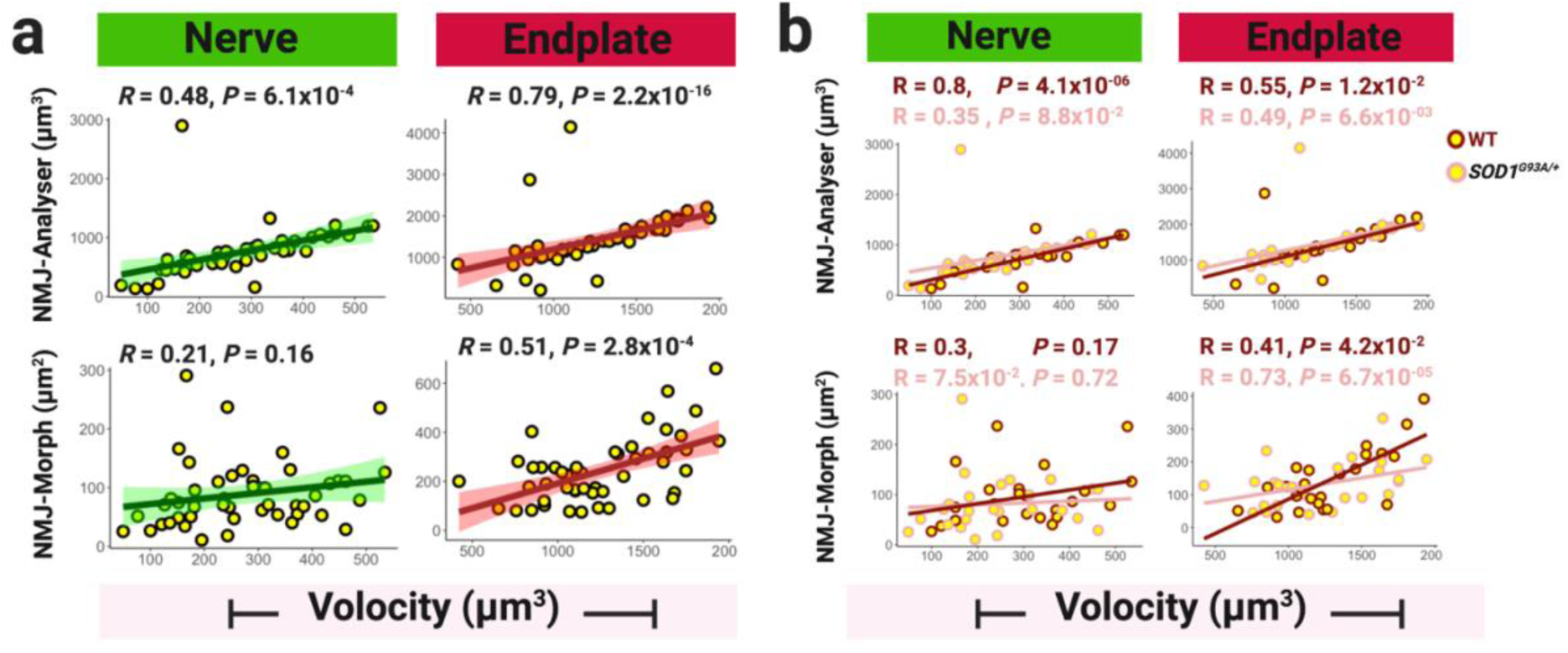
Performance of NMJ-Analyser and NMJ-Morph using batch 2. (a) Correlation of nerve terminal and endplate output obtained using NMJ-Analyser and NMJ-Morph. Datapoint showed non-normal distribution and spearman correlation tests were used. Green and red shading represent the confidence interval. (c) Correlation of nerve terminal and endplate output obtained using NMJ-Analyser and NMJ-Morph, divided by genotype.

**Table S1.**
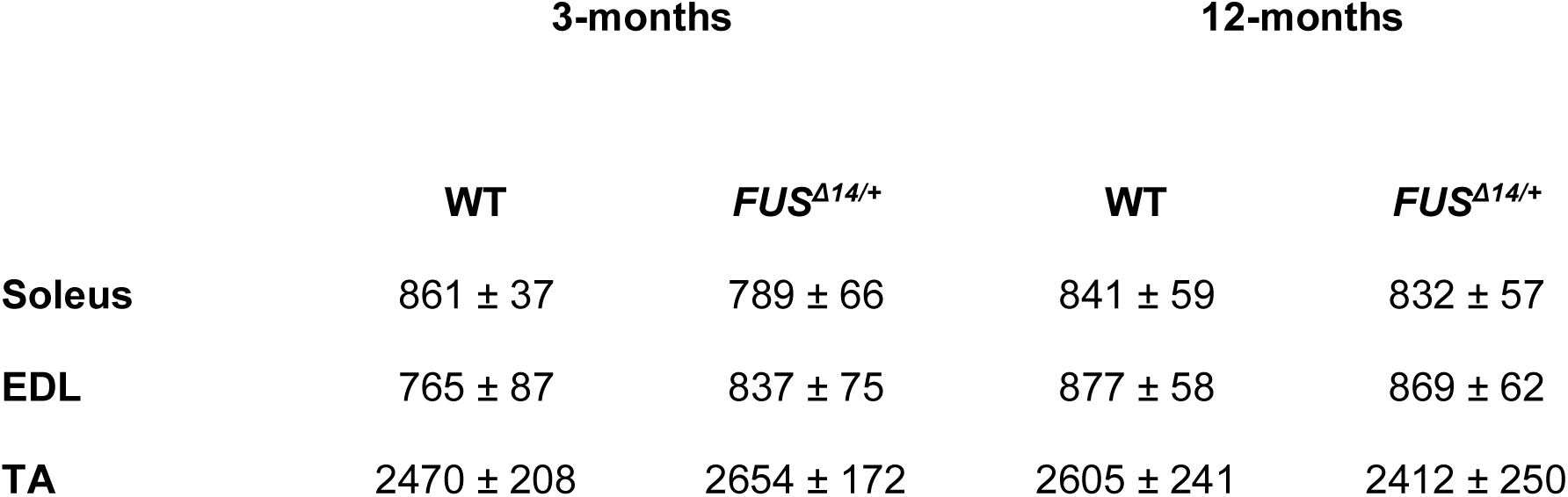
Total number of fibres in hindlimb muscles.

|  | 3-months |  | 12-months |  |
| --- | --- | --- | --- | --- |
|  | WT | *FUS*<sup>Δ14/+</sup> | WT | *FUS*<sup>Δ14/+</sup> |
| **Soleus** | 861 ± 37 | 789 ± 66 | 841 ± 59 | 832 ± 57 |
| **EDL** | 765 ± 87 | 837 ± 75 | 877 ± 58 | 869 ± 62 |
| **TA** | 2470 ± 208 | 2654 ± 172 | 2605 ± 241 | 2412 ± 250 |

**Table S2.**
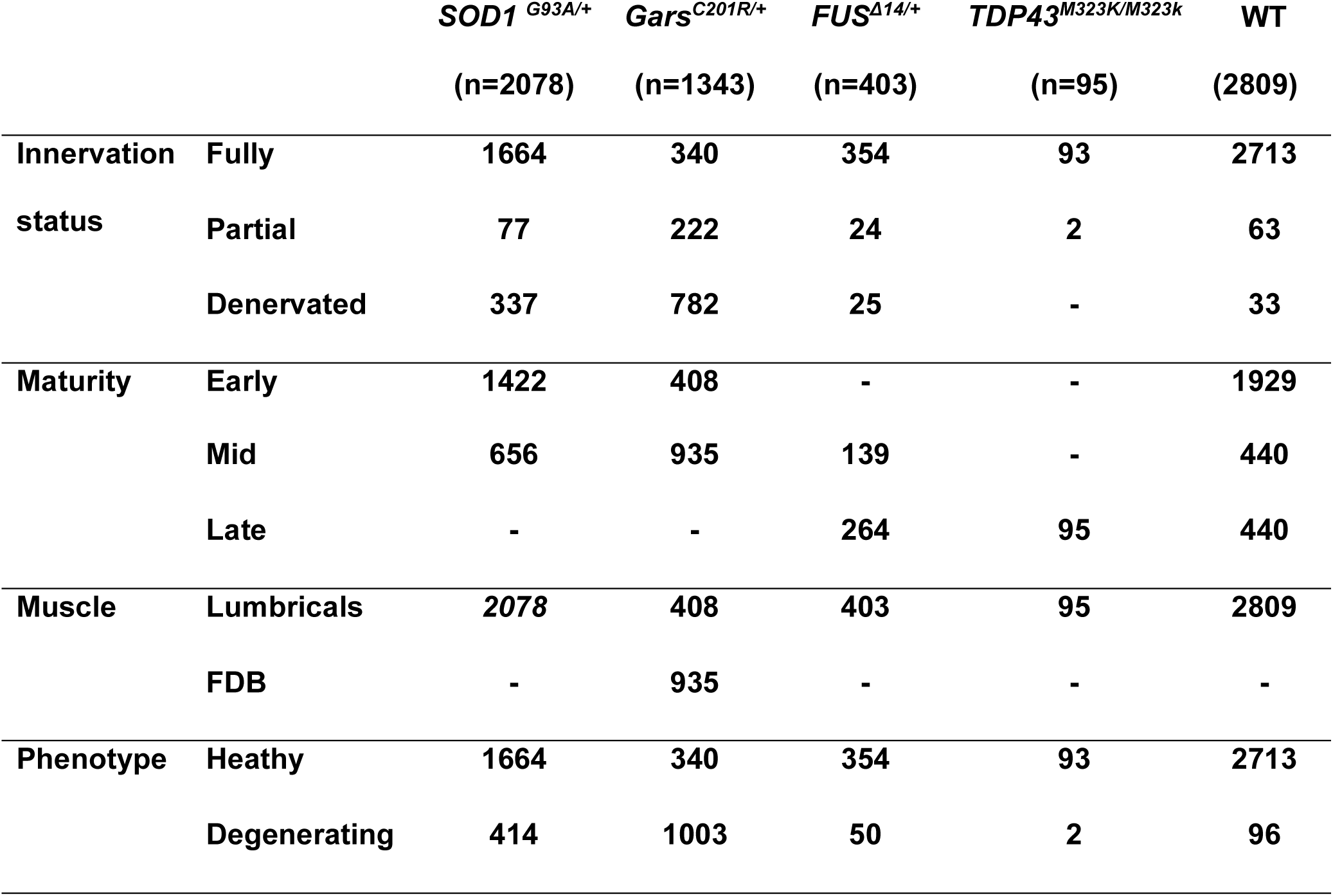
Total number of NMJs analysed.

## Additional File 2

### NMJ-Analyser Tutorial

#### Running NMJ-Analyser

The NMJ-Analyser is a pipeline originally written in Python language.

##### Step 1

- Confirm if Python is installed in your machine. Mac OSX: type ‘terminal’ in spotlight search. Type ‘python’ to confirm it is installed. A line command pops up. Windows: type ‘cmd’ in Start menu. Type ‘python --version’ to confirm it is installed. A line command pops up.
- If Python is not installed, go to https://www.python.org/downloads/

#### Step 2

To run NMJ-Analyser, need to install the following modules:

- SciPy, for multidimensional image processing: https://www.scipy.org/install.html
- PIL, for reading images: https://pillow.readthedocs.io/en/stable/installation.html
- Glob, for finding the pathnames: https://docs.python.org/3/library/glob.html
- OS: https://pypi.org/project/os-sys/
- Numpy: https://numpy.org/install/
- Argparse: https://pypi.org/project/argparse/
- Pandas: https://pandas.pydata.org/pandas-docs/stable/getting_started/install.html
- Sys: https://docs.python.org/3/install/

##### Step 3

- 3D images should be converted into individual nerve terminal and endplate files (plane view, .TIFF,.JPEG or .PNG).
- Create a folder ‘MouseTest’ with images file containing the keyword red or RED (endplate staining), or green or GREEN (nerve staining). The images should be ordered numerically (*i.e.* Mouse1_GREEN_0001.jpg…. Mouse1_GREEN_0010.jpg).
- Identify the directory of ‘Test’ folder and Processing.py file. /Users/’name_user’/…/folder…
- Alternatively, download the ‘MouseTest’ folder (sample images for training) from https://github.com/csudre/NMJ_Analyser.

##### Step 4

- Download NMJ-Analyser scripts from https://github.com/csudre/NMJ_Analyser. Find the ProcessingGARS.py file.
- Set up the directory (folder where you store the working files) and path files.
- Line 45 in the ProcessingGARS.py file: the plane of resolution (size of pixel, in yellow) and slice thickness (in red).
- Line 45 in the ProcessingGARS.py file: minimum size of signal to be considered as

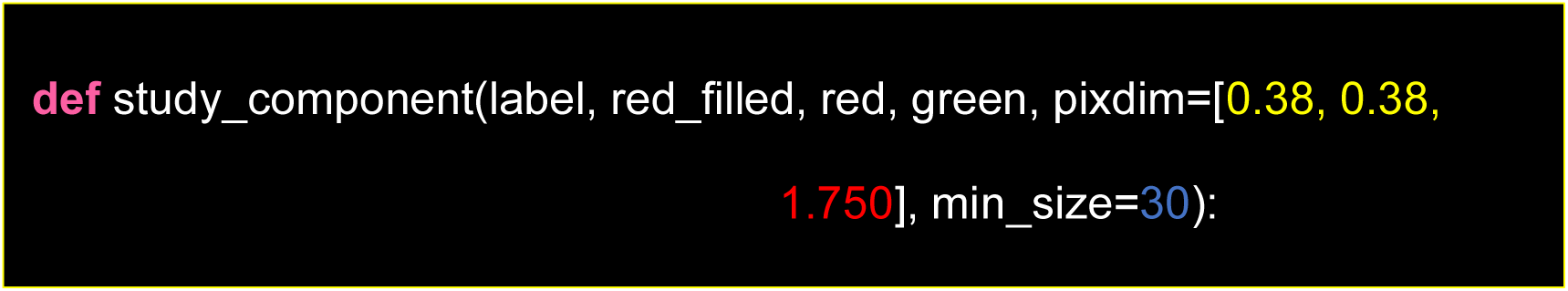
- Line 215: default threshold is established as 64.

##### Step 6

- In the terminal, type the following: (in yellow =pixel size, in red=slice thickness)
- python /’your directory’ /NMJ_Analyser-master/ProcessingGARS.py -p /’your directory’ /MouseTest -dx 0.277 -dz 1.5.
- A .csv file containing a number of parameters will pop. Morphological features for each nerve terminal (green) and endplate (red).

#### Normalization

Batch effects are non-biological variation between experiments performed at different timepoints [48,49]. Variables contributing to batch effect (for example, fixation, penetration of antibodies, image thresholding and background staining) impact on the quality and reliability of the immunofluorescence staining. NMJ-Analyser considers the variability between batches by using the same thresholding protocol for all sections and using the voxel size as an input to calculate the 3D NMJ features. We considered a minimum and maximum cut-off size of NMJ structures that are automatically included in the pipeline (Additional File 1: Fig.S1). In cases where multiple batches were compared (i.e. for C57BL/6J and C57BL/6J-SJL male mice), the following normalization protocol was applied: 1) use the same thresholding value and background correction across batches, 2) compare the mean fluorescence intensity (MFI) of nerve terminal and endplate in WT samples, 3) use the cumulative distribution function (ECDF) and Kolmogorov-Smirnov test as a guide of MFI differences across samples (if not normally distributed), and 4) divide the value of each pre- and post-synaptic morphological variable by their MFI. Details of the normalization protocol and R/RStudio scripts can be found at Additional File 1: Fig.S1.

Normalization is required when multiples batches are analysed together. Normalization procedure assumes the imaging setup was maintained constant, except for the pixel size.

##### Step 1

- Maintain the same thresholding value and background correction across batches

##### Step 2

- Only in wildtype samples, identify whether the nerve terminal or endplate MFI follows normal distribution (Shapiro-Wilk test, a,b).
- Compare mean fluorescence intensity (MFI) of nerve terminal and endplate in WT samples of each batch. use the cumulative distribution function (ECDF) and Kolmogorov Smirnov test or Kruskal-Wallis test as a guide of MFI differences across samples If significantly differ go to step 3 (Fig.S1).

**Fig.S1.**
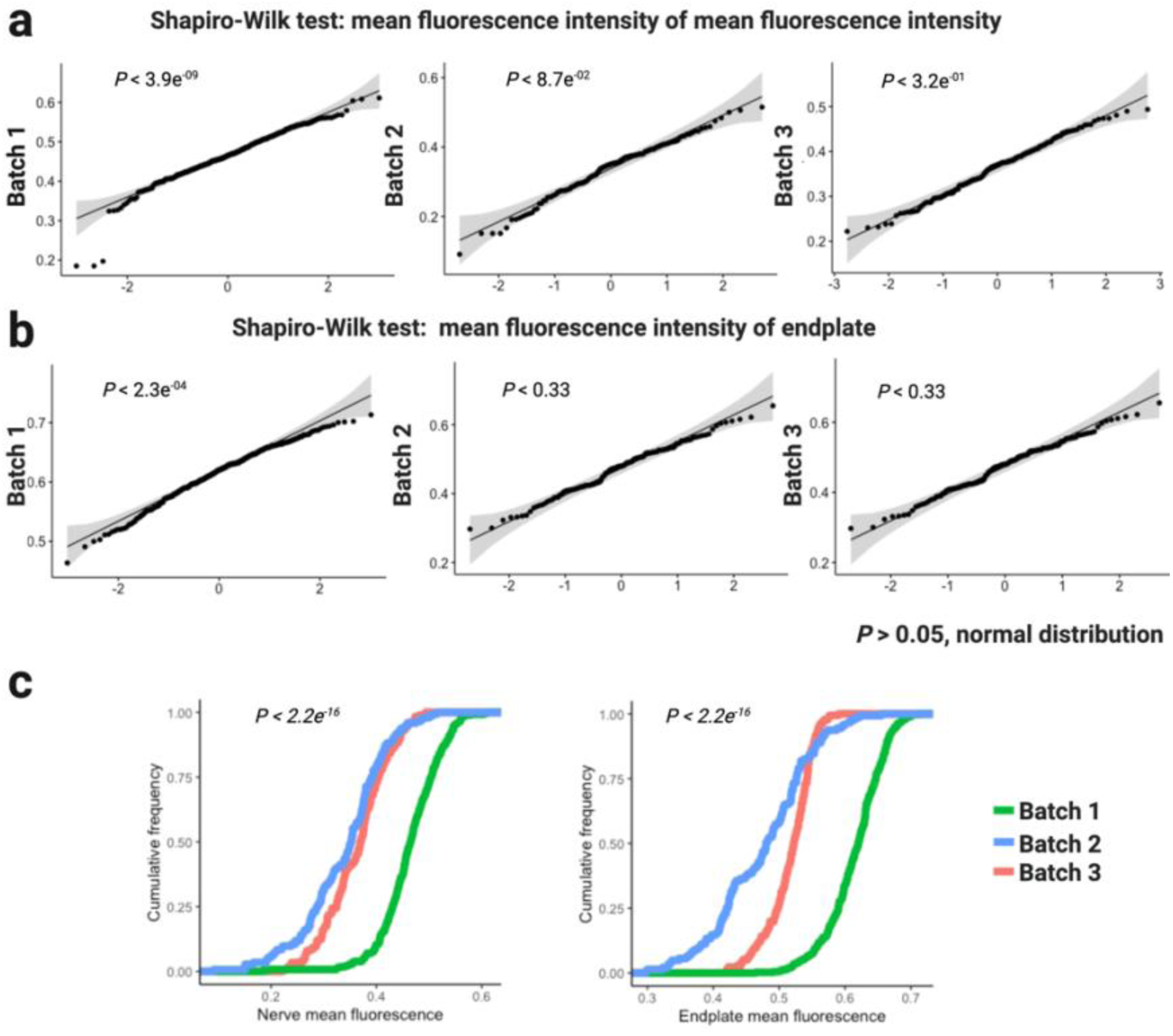

##### Step 3

- Divide the value of each pre- and post-synaptic morphological variable by their corresponding MFI.

#### Manual curation

Images containing multiple NMJs require manual inspection. This procedure is required to fully identify individual NMJs and avoid poor counting. Thus, it is possible that two or more NMJs can be counted as one if they are close (<20μm).

##### Step1

- Download and install ITK-SNAP viewer: http://www.itksnap.org/pmwiki/pmwiki.php
- Open the confocal image (.lsm5 format) on a ITK-SNAP.
- Identify each NMJ obtained from NMJ-Analyser .csv output by typing the position (x,y,z) on ‘cursor position’ (Fig.S2).

**Fig.S2.**
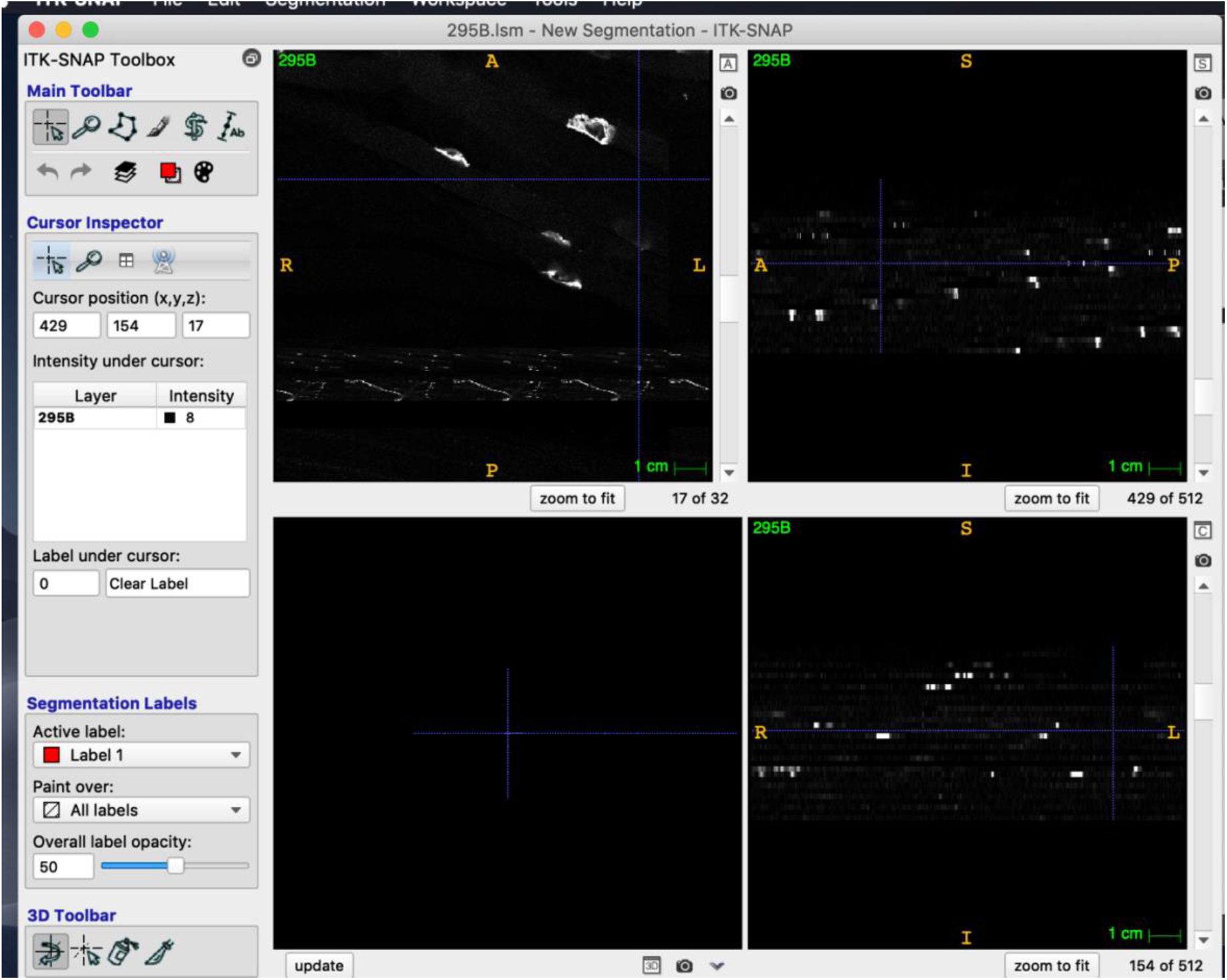

#### Machine Learning

Machine learning pipeline and instructions to run the machine learning platform can be downloaded from https://github.com/SethMagnusJarvis/NMJMachineLearning

#### NMJ-Analyser validation

We compared the nerve terminal and endplate volume outputs obtained by of NMJ-Analyser and Volocity. The figure below shows strong correlation between outputs obtained by NMJ-Analyser and Volocity (Fig.S3)

**Fig.S3.**
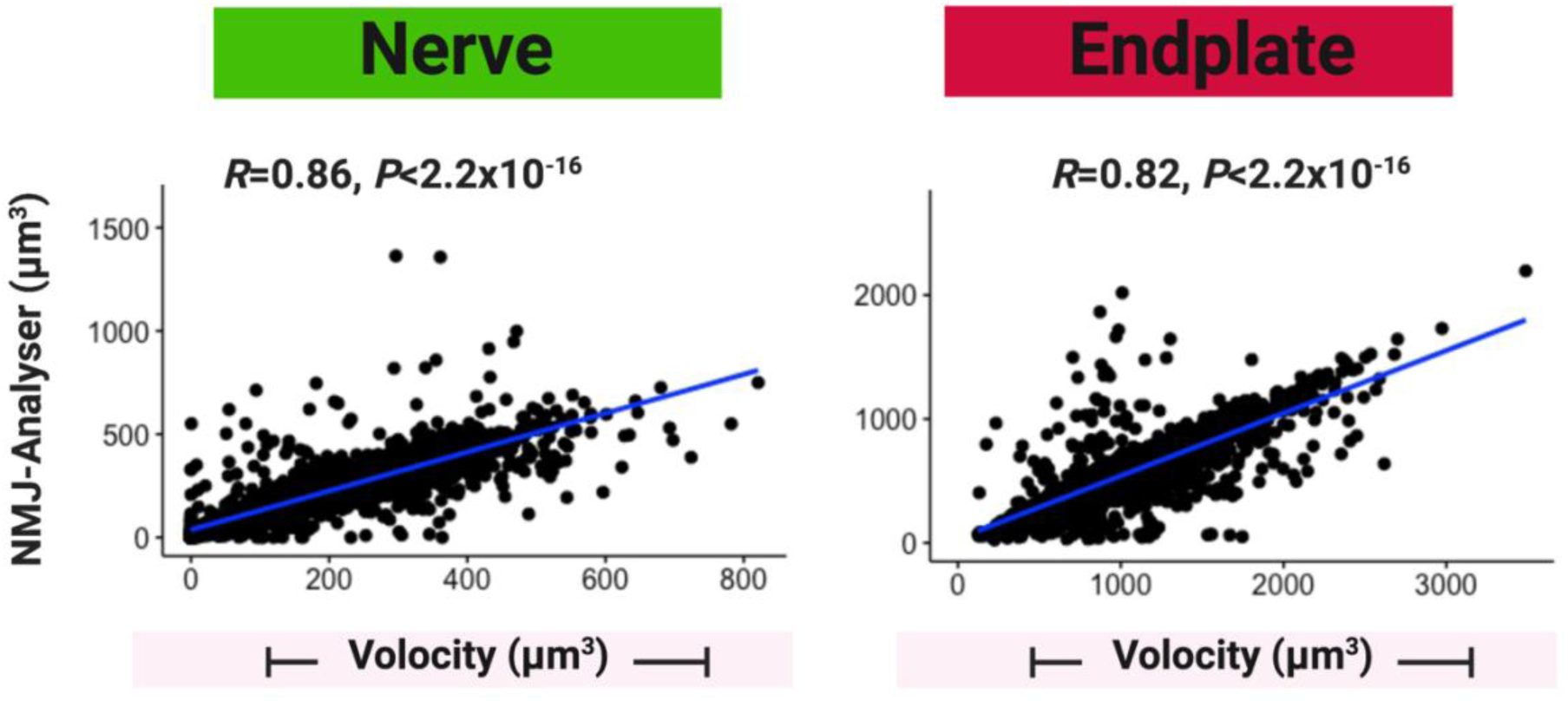

